# Transcription factor binding process is the primary driver of noise in gene expression

**DOI:** 10.1101/2020.07.27.222596

**Authors:** Lavisha Parab, Sampriti Pal, Riddhiman Dhar

## Abstract

Noise in expression of individual genes gives rise to variations in activity of cellular pathways and generates heterogeneity in cellular phenotypes. Phenotypic heterogeneity has important implications for antibiotic persistence, mutation penetrance, cancer growth and therapy resistance. Specific molecular features such as presence of the TATA box sequence and promoter nucleosome occupancy have been associated with noise. However, the relative importance of these features in noise regulation has not yet been assessed. In addition, how well these features can predict noise also remains unclear. Here through an integrated statistical model of gene expression noise in yeast we found that the number of regulating transcription factors (TFs) was a key predictor of noise. With an increase in the number of regulatory TFs, we observed a rise in the number of TFs that bound cooperatively. In addition, increased number of TFs meant more overlaps in TF binding sites that caused competition between TFs for binding to the same region of the promoter. Through modeling of TF binding to promoter and applying stochastic simulations, we demonstrated that competition and cooperation among TFs could increase noise. Thus, our work uncovers a process of noise regulation that arises out of the dynamics of gene regulation and is not dependent on any specific transcription factor or specific promoter sequence.

## Introduction

Random fluctuations in molecular events operating inside a cell generate variations in the expression levels of genes that is termed as gene expression noise. Expression noise gives rise to variations in activities of cellular pathways and generates phenotypic heterogeneity among individual cells of an isogenic population under identical environmental condition. Gene expression noise has important role in antibiotic persistence (Balaban *et al*., 2004; Rotem *et al*., 2010; Wakamoto *et al*., 2013; Arnoldini *et al,* 2014; Page and Peti, 2016) and incomplete penetrace of mutations (Eldar *et al*., 2009; Raj *et al*., 2010; Burga *et al*., 2011; Dickinson *et al*., 2016; Taeubner *et al*., 2018). In addition, phenotypic heterogeneity has a key role in growth of cancers (Meacham and Morrison, 2013; Nguyen *et al*., 2019; Sharma *et al*., 2019) and in emergence of therapy resistance (Shaffer *et al*., 2017; Hammerlindl and Schaider, 2018; Gupta *et al*., 2019; Farquhar *et al*., 2019; Emert *et al*., 2021).

Gene expression noise has been measured in some microbial systems (Newman *et al*., 2006; Bar-Even, A. *et al*., 2006; Taniguchi *et al*., 2010; Silander *et al*., 2013) and its molecular origins have been widely investigated (McAdams and Arkin, 1997; Elowitz *et al*., 2002; Blake *et al*., 2003; Raser and O’Shea, 2004; das Neves *et al*., 2010; Sanchez *et al*., 2011; Hornung *et al*., 2012; Salari *et al*., 2012; Sanchez *et al*., 2013; Sharon *et al*., 2014; Chen and Zhang, 2016; Wu *et al*., 2017; Faure *et al*., 2017; Baudrimont *et al*., 2019). These studies have shown a correlation between presence of the TATA box motif in the promoter region of a gene and expression noise (Bar-Even *et al*., 2006; Tirosh *et al*., 2006; Hornung *et al*., 2012; Ravarani *et al*., 2016). Further, promoter nucleosome occupancy, alone as well as in combination with presence of the TATA box motif, and histone modification patterns have also been associated with expression noise (Tirosh and Barkai, 2008; Choi and Kim, 2009; Small *et al*., 2014; Weinberger *et al*., 2012; Chen and Zhang, 2016; Faure *et al*., 2017; Nicolas *et al*., 2018). These features can influence transcriptional burst size and frequency (Donovan *et al*., 2019; Larsson *et al*., 2019; Nicolas *et al*., 2018) which, in turn, impact expression noise (Hornung *et al*., 2012; Zoller *et al*., 2015; Wang *et al*., 2019; Engl *et al*., 2020). However, even after so many studies over the years, the relative importance of these molecular features in noise regulation remains unknown. In addition, to what extent each of these molecular features can predict noise has not yet been quantified. A predictive model of noise will be immensely helpful for a better understanding of regulation of expression noise in biological systems.

In the current work, we report development of an integrated statistical model of gene expression noise in yeast by combining a large number of molecular features that can impact gene expression. We quantified the relative contribution of each of these features in explaining variations in noise values of genes and tested their predictive abilities. We uncovered that the number of regulatory TFs of a gene was an important predictor of noise. Increase in the number of regulatory TFs was associated with a concomitant increase in the number of cooperative TFs. In addition, increase in the number of regulatory TFs meant crowding of TF binding sites in the promoter region of a gene. This led to more overlaps between TF binding sites, thereby increasing competition between TFs for binding to the same promoter site. Mathematical modeling and stochastic simulations showed that a mere increase in the number of TFs could not explain the increase in expression noise, whereas cooperative and competitive TF binding could lead to higher expression noise. Taken together, our work demonstrates that the binding process of transcription factors is the best predictor of noise in yeast. Our work uncovers a dynamic noise regulation mechanism originating from competition and cooperation among transcription factors. This mechanism is not dependent on specific transcription factor or specific promoter sequence and thus, could be of interest to researchers working on different biological organisms.

## Results

### Quantification of expression noise at the level of mRNA and protein

We quantified gene expression noise at the level of both mRNA and protein using two different experimental datasets. For calculating noise at the level of mRNA, we used single-cell RNA-seq data in yeast from Nadal-Ribelles *et al*., 2019 (Fig. 1A). The dataset contained expression values of genes in 127 single-cells of *Saccharomyces cerevisiae* strain BY4741 grown in rich growth medium (YPD) and expression profiles measured at early-log phase. We obtained expression values of 5475 genes from this dataset. To quantify noise, we used a measure of noise that was independent of mean expression level through fitting a spline to the noise (coefficient of variation, CV) vs mean plot and calculating vertical distance of noise values from the fitted curve (Fig. 1A; Supplementary fig. S1). We obtained noise values at the protein level for 2763 genes in *S. cerevisiae* S288C strain grown in rich medium (YPD) (Newman *et al*., 2006; Fig. 1B). We used their measure of distance to median (DM) as the measure of noise in our study (Supplementary fig. S1). Noise at the mRNA level showed significant correlation with noise at the protein level (r=0.44, p=1.7×10^-95^ Fig. 1C) although the range of absolute noise values were very different. Genes showed a wide range of expression noise values with highly noisy genes showing high positive values and low noise genes showing large negative values.

To quantify the relative importance of each molecular feature in noise regulation and to measure their ability to predict noise, we randomly segregated the noise data into training (80% of the full data) and test datasets (remaining 20%) (Fig. 1D). For quantifying predictive ability of a single feature, we fitted a linear regression model to the training data at this step. For quantifying predictive ability of a combination of features, we first removed multicollinear features and identified the key set of features through Ridge or Lasso regression on the full data and then fitted a linear regression model on the training data. This gave us the fraction of variation explained by the model (Fig. 1D). We used the fitted model to make predictions on the test data and computed predicted R^2^ values (Fig. 1D). We performed this analysis in both mRNA and protein noise datasets to ensure that inferences drawn were not biased by a specific dataset.

Molecular features that had been associated with expression noise, such as presence of the TATA box sequence in the promoter (Bar-Even *et al*., 2006; Tirosh *et al*., 2006; Hornung *et al*., 2012; Ravarani *et al*., 2016) and promoter nucleosome occupancy (Tirosh and Barkai, 2008; Choi and Kim, 2009; Small *et al*., 2014) showed association with noise (Supplementary fig. S2) but were poor predictors (Fig. 1E). Specifically, the TATA box sequence, promoter nucleosome occupancy alone and in conjunction with the TATA box sequence could explain only ~2-4%, ~6-7% and ~8-9% of the noise variation, respectively and had low predictive power (predicted R^2^ value 0.02-0.03 for the TATA box alone, 0.05-0.07 for promoter nucleosome occupancy alone, and 0.07 for the TATA box + promoter nucleosome occupancy; Fig. 1E). This suggested that these features were largely associated with noise and were not predictive.

**Figure 1.**
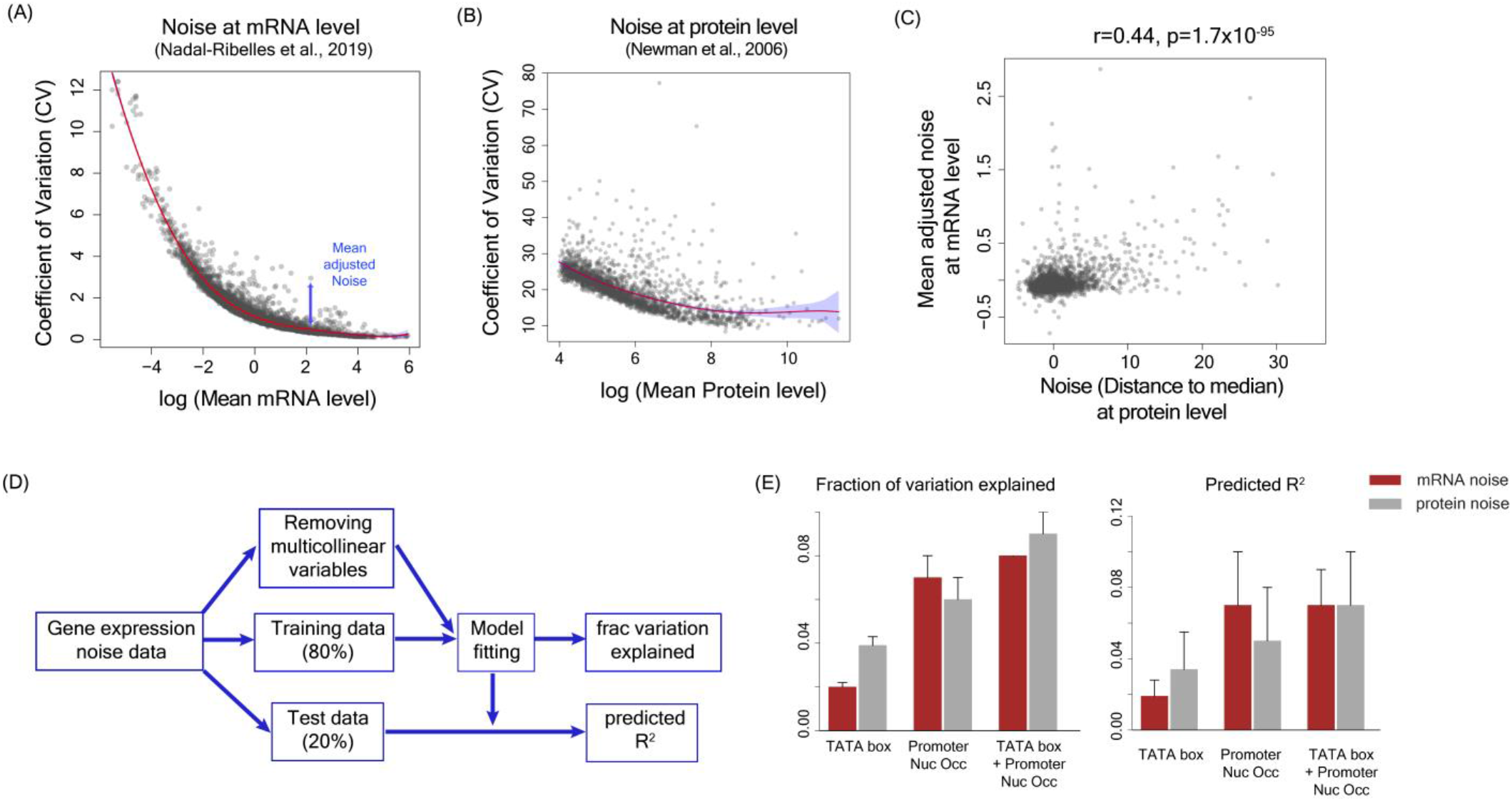
Presence of the TATAbox sequence and promoter nucleosome occupancy are poor predictors of gene expression noise. **(A)** Noise values calculated at the mRNA level from single cell RNA-seq data in yeast (Nadal-Ribelles *et al*., 2019). The mean adjusted noise was calculated by fitting a polynomial curve to the CV vs mean plot shown by the red line. Each point shows CV and mean mRNA level values for a gene. **(B)** Noise values calculated at the protein level from flow cytometry measurements by Newman *et al*., 2006. The red line shows the best polynomial fit and the shaded blue region shows 95% confidence interval. **(C)** Noise at the mRNA level was significantly correlated with noise at the protein level (r=0.44, p=1.7×10^-95^ with the null hypothesis that the correlation coefficient is zero). **(D)** Flowchart showing the steps for model fitting, calculation of fraction of variation explained and derivation of predicted R^2^. **(E)** Fraction of variation explained and predicted R^2^ values by presence or absence of the TATA box sequence, average promoter nucleosome occupancy per nucleosome bound site and the combination of presence or absence of the TATA box sequence with promoter nucleosome occupancy.

### An integrated statistical model of noise for evaluating contribution of all molecular features

To identify molecular features that could explain the observed variations in noise values and predict noise, we built an integrated statistical model considering all features that were known or were likely to influence gene expression, as these could be potential regulators of noise (Fig. 2, Supplementary table S1). The goals of the integrated statistical model were to test the predictive power of each molecular feature individually and to identify the best sets of features for noise prediction out of a large number of possible combinations.

**Figure 2.**
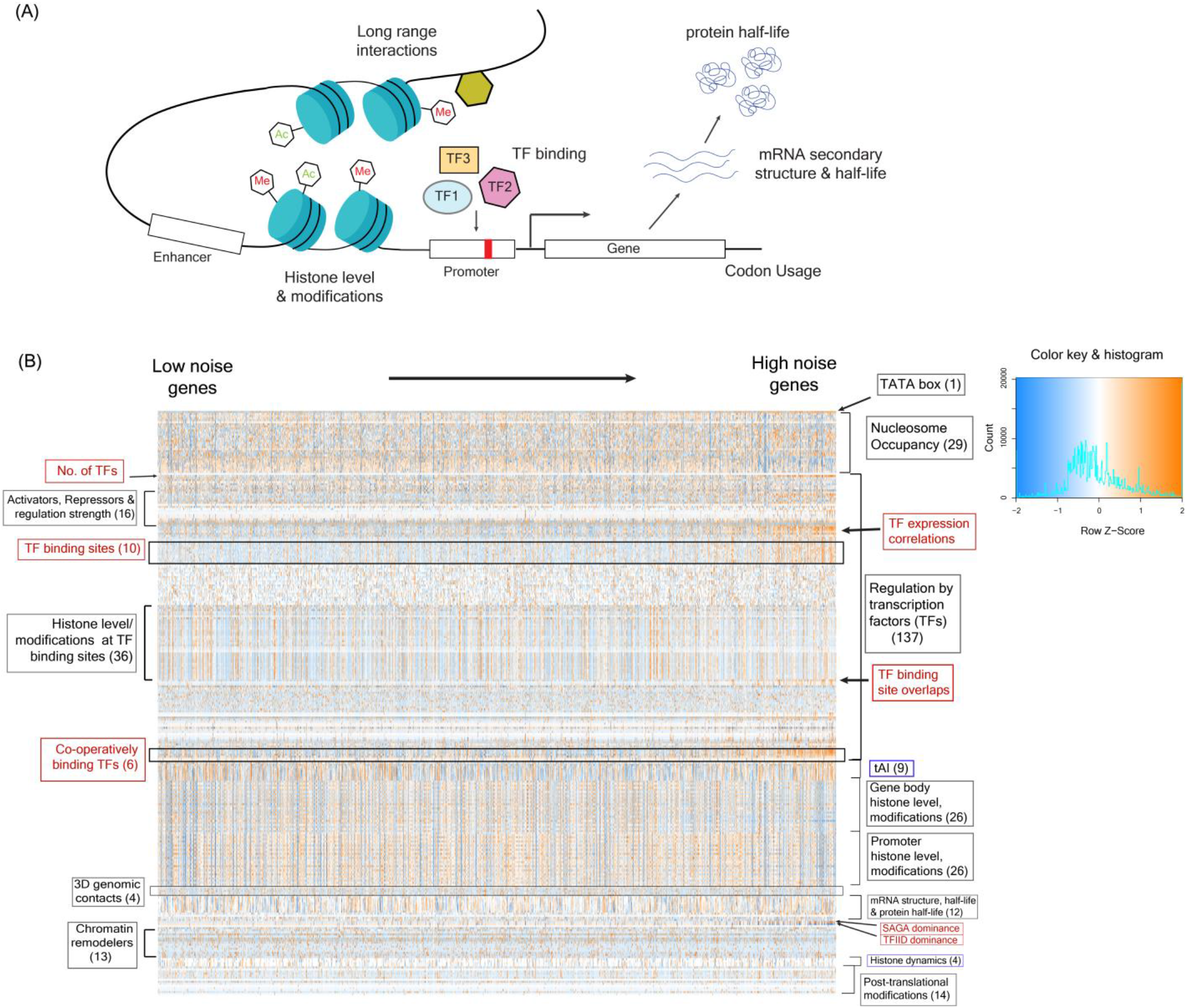
An integrated statistical model of gene expression noise. **(A)** Schematic diagram depicting the molecular features that could impact gene expression and thus, could have a key role in regulation of expression noise. **(B)** An integrated model of noise constructed considering the TATA box sequence, absolute nucleosome occupancy levels, gene regulation by TFs, tRNA adaptation index, histone modification pattern in gene-body and promoter regions, 3D genomic contacts, mRNA structure and half-life, protein half-life, activity of chromatin remodelers, histone binding dynamics and post-translational modifications. The heatmap shows values of all these features (scaled and centered) in genes (represented in the columns) sorted according to their noise values at the protein level. The number of features for which the data is shown in the heatmap are indicated inside the brackets. Features highlighted in red appear different in their values between low and high noise genes. The panel on the right shows the color key for the heatmap along with the distribution of values of all features (histogram).

The molecular features incorporated in the integrated model included the number of regulating TFs, location of their binding sites, their mean expression and noise levels, SAGA/TFIID dependence of genes for their expression (Huisinga and Pugh, 2004), whether a gene was co-activator redundant or TFIID dependent (Donczew *et al*., 2020), binding activity of several broadly acting TFs such as TBP, ABF1 and RAP1 (van Werven *et al*., 2009; Lickwar *et al*., 2012; de Jonge *et al*., 2020), binding patterns of chromatin remodelers (Yen *et al*., 2012; Zentner and Henikoff, 2013; Ramachandran *et al*., 2015), histone levels, histone modification patterns and histone binding dynamics (Pokholok *et al*., 2006; Dion et al., 2007), three-dimensional genomic contacts (Duan *et al*., 2010), tRNA adaptation index (Tuller *et al*., 2010), mRNA secondary structure, mRNA and protein half-lives (Kertesz *et al*., 2010; Geisberg *et al*., 2014; Belle *et al*., 2006), post-translational modifications (Ledesma *et al*., 2018), in addition to nucleosome occupancy pattern (Oberbeckmann *et al*., 2019) and presence/absence of the TATA box sequence in the promoter (Rhee and Pugh, 2012). For a gene we only considered those TFs for which experimental evidence for DNA binding had been obtained or change in expression upon knocking out the TF had been experimentally observed. For nucleosome occupancy, we not only considered the number of nucleosome-bound sites but included the absolute nucleosome occupancy pattern (Oberbeckmann *et al*., 2019). In total, we considered 329 features in our integrated model (Supplementary table S1).

### Molecular features associated with TF binding were the top predictors of noise

We tested each feature individually for its ability to explain variation in the noise data and to predict noise in both mRNA and protein noise datasets. We then ranked these features according to the fraction of variation explained and by predicted R^2^ values. The ranking of the features, whether based on fraction of variation explained or predicted R^2^ value were substantially correlated among mRNA and protein noise datasets with correlation values of 0.67 and 0.76 respectively (Fig. 3A, B).

**Figure 3.**
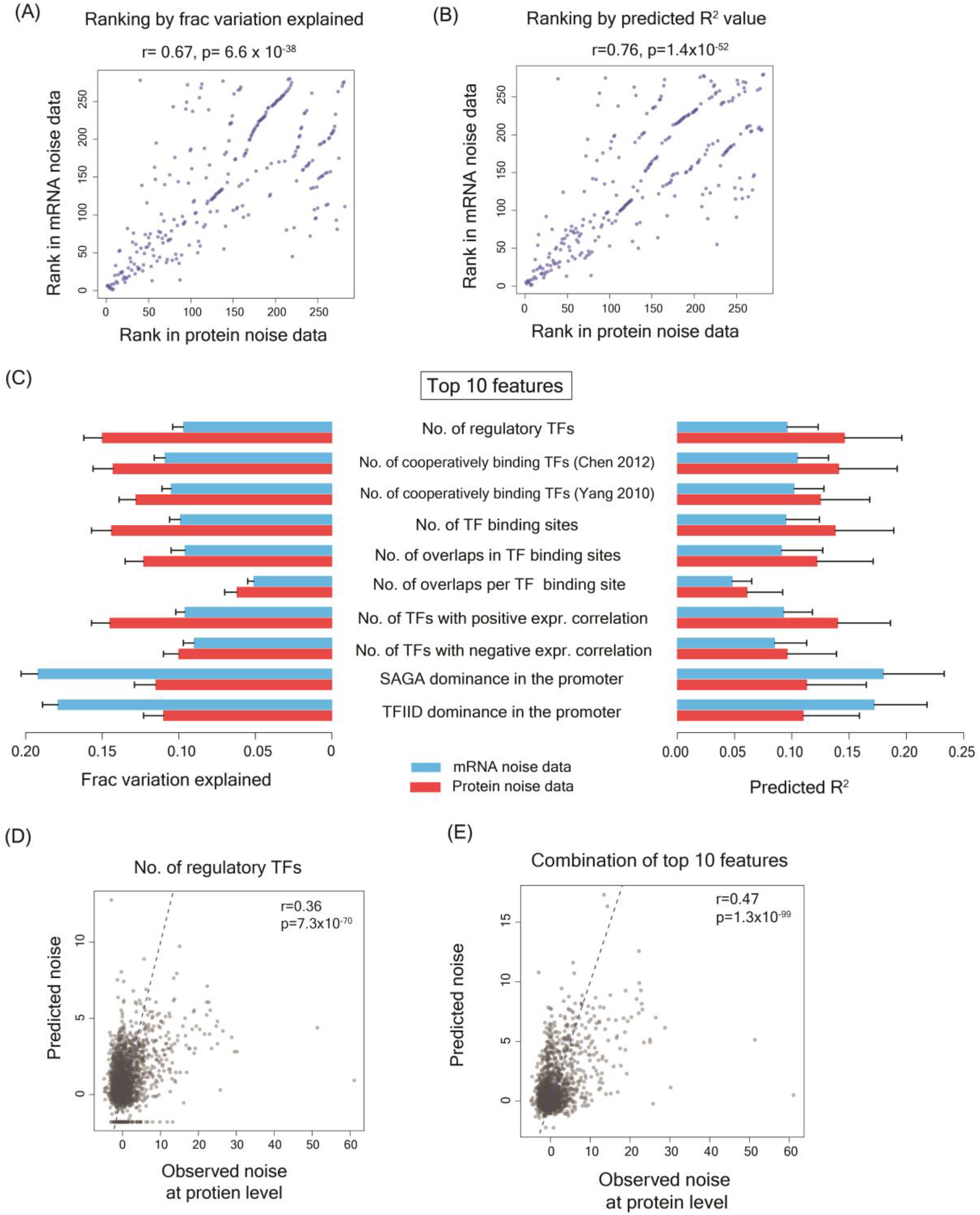
Features with highest predictive powers were largely related to transcription factor binding process. **(A)** Rankings of features according to the fraction of variation explained in mRNA noise dataset and in protein noise dataset were highly correlated (r=0.67, p=6.6×10^-38^ with null hypothesis that the correlation value is zero) **(B)** Rankings of features according to the predicted R^2^ value in mRNA and protein noise datasets were highly correlated (r=0.76, p=1.4×10^-52^ with the null hypothesis that the correlation value is zero) **(C)** Fraction of variation explained and predicted R^2^ value for top 10 features for both mRNA and protein noise datasets. **(D-E)** Observed and predicted noise values by linear regression model considering a single feature (number of regulatory TFs) (D) and by the combination of top 10 features (E).

The top 10 features for explaining the variation existing in the noise data and for predicting noise values contained the same features although their rankings were slightly different. The distributions of values of some of these features are shown in supplementary fig. S3. Interestingly, eight of these features were associated with TF binding, suggesting a key role for TFs in noise regulation (Fig. 3C). These included number of regulatory TFs for a gene (fraction of variation explained ~0.1-0.15 and a predicted R^2^ of ~0.1-0.15) and the number of TF binding sites (fraction of variation explained ~0.1-0.14, predicted R^2^ ~0.1-0.14). The remaining two features were related to SAGA-dependence and TFIID-dependence of genes for their transcription. Stress response genes in yeast are known to be noisier than housekeeping genes (Newman *et al*., 2006). While housekeeping genes are dependent on TFIID complex for their expression, stress response genes are usually SAGA complex dependent. SAGA dependence and TFIID dependence could explain 0.11-0.19 and 0.11-0.17 fraction of variation respectively with predicted R^2^ values of 0.11-0.18 and 0.11-0.17 respectively (Fig. 3C).

We further validated predictive abilities of these features by correlating the observed and predicted noise values. Predicted values obtained using number of regulatory TFs as the only feature and the combination of top 10 features showed correlations of 0.36 (p=7.3×10^-70^ with the null hypothesis that the correlation value is zero) and 0.47 (p=1.3×10^-99^) respectively with the observed noise values at the protein level (Fig. 3D, 3E).

Of all the features in our model, TF binding process could explain the largest part of the faction of variation in the data and had the highest predictive power (Fig. 4). The integrated model comprising of all features was able to explain 0.46 fraction of the variation in noise at the mRNA level and 0.47 fraction of the variation in noise at the protein level (Fig. 4A). TF binding alone explained 0.26 fraction of the variation in the noise at the mRNA level and 0.30 fraction of the noise at the protein level (Fig. 4A). In addition, the integrated model was able to predict noise at the mRNA level with predicted R^2^ value of 0.31 and at the protein level with predicted R^2^ value of 0.36 (Fig. 4B). As before, TF binding process alone could predict noise at both mRNA and protein levels with predicted R^2^ value of 0.23 (Fig. 4B).

**Figure 4:**
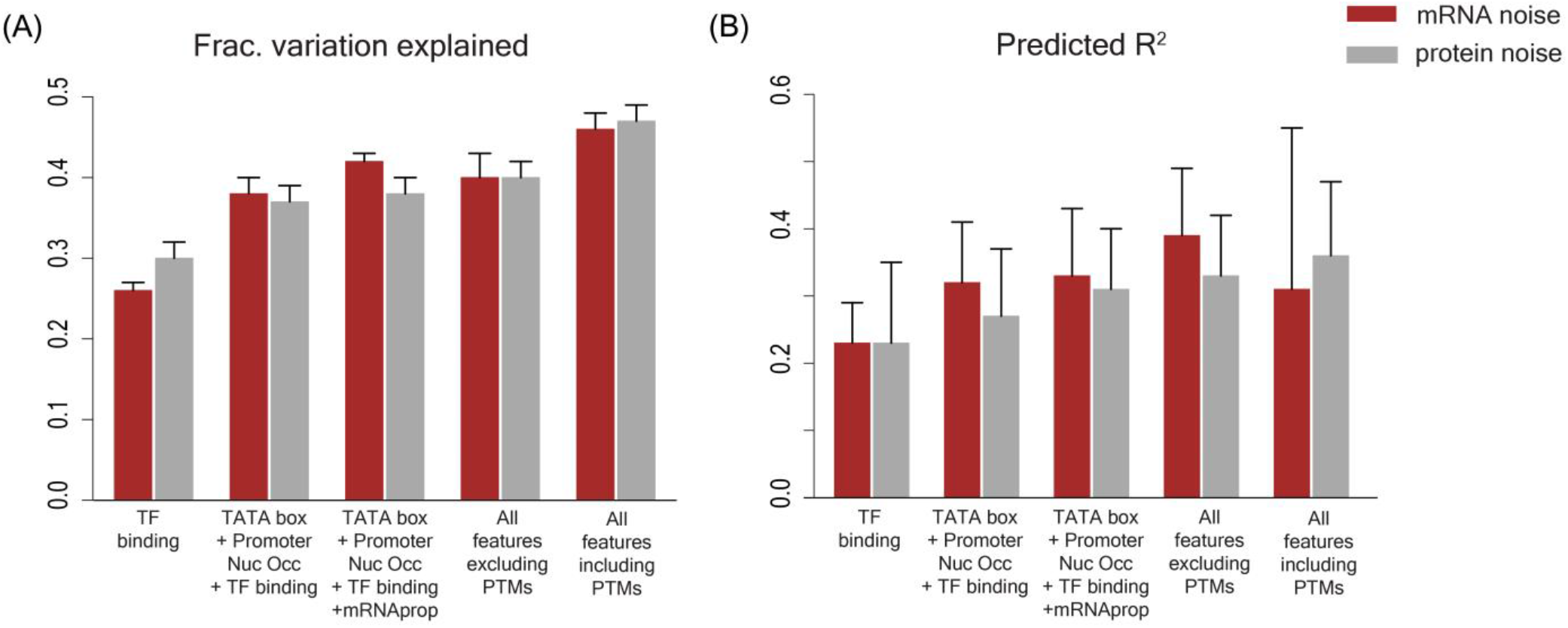
**(A)** Fraction of variation explained in gene expression noise data and **(B)** predictive ability (given by predicted R^2^ value) by features associated with TF binding; combination of TF binding with the TATA box sequence and promoter nucleosome occupancy; combination of TF binding with TATAbox, promoter nucleosome occupancy and mRNA properties; combination of all features excluding post-translational modifications (PTMs), and by combination of all features including PTMs.

Several genes in the yeast genome have been retained from a whole-genome duplication (Byrne and Wolfe, 2005) and thus share many molecular features including promoter and coding region sequences with their duplicates. This could bias our analysis and can lead to inflated predictive R^2^ values. Thus, to assess the impact of gene duplicates on our analysis, we removed duplicates from our datasets and repeated all analysis. The fraction of variation explained and predicted R^2^ values by individual features and by combinations of features were comparable between datasets with and without duplicate genes (Supplementary figs S4, S5).

### Genes with high expression noise were regulated by a higher number of TFs

Our model revealed a significant correlation between the number of regulating TFs of a gene and noise, at both mRNA and protein levels (for protein noise r=0.36, p=7.3×10^-70^; Fig. 5A; for mRNA noise r=0.26, p=5.7×10^-85^; Supplementary fig. S6A). We further classified genes into 20 bins sorted according to their noise values equally spaced noise bins barring the first and the last bins. The first bin had open-ended lower limit for noise values to include genes showing very low noise levels. The last bin had open-ended upper limit for noise values so as to include genes showing very high noise levels. This helped us avoid having bins with a very low number of genes. We then looked at the distribution of the number of regulatory TFs of genes in these bins (Fig. 5B, Supplementary fig. S6B). The genes in the highest noise bins on average had >75% more regulatory TFs compared to the genes in the lowest noise bins (Fig. 5B, Supplementary fig. S6B).

**Figure 5.**
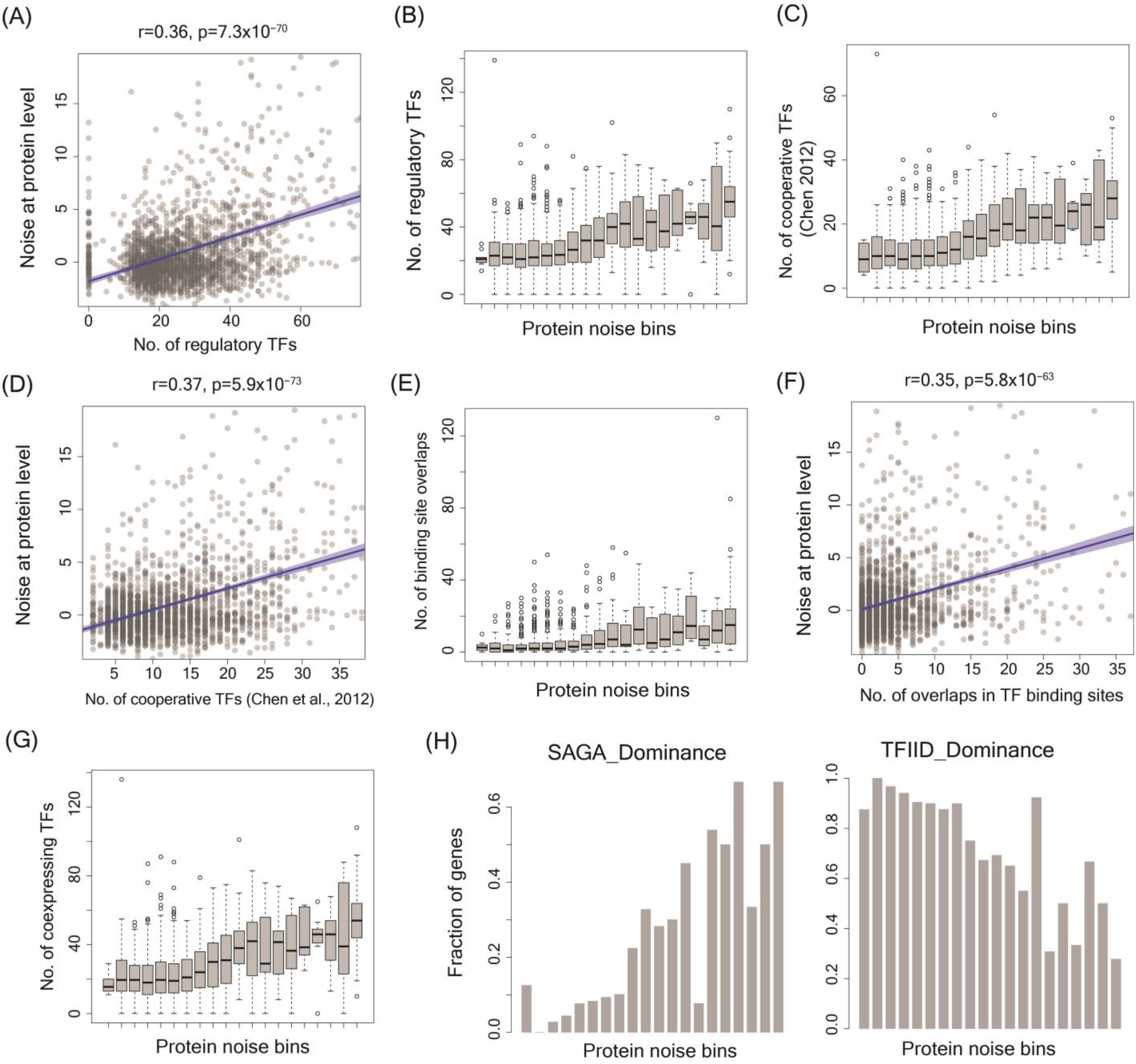
Genes with high noise were regulated by a higher number of TFs, had a higher number of cooperatively binding TFs, and showed more overlaps in TF binding sites compared to low-noise genes. **(A)** Correlation between noise at protein level and the number of regulatory TFs. **(B)** Number of regulatory TFs of genes across 20 protein noise bins. **(C)** Number of cooperative TFs of genes across protein noise bins. **(D)** Correlation between noise at protein level and the number of cooperative TFs (Chen et al., 2012) **(E)** Number of overlaps between TF binding sites for genes across protein noise bins. **(F)** Correlation between noise at protein level and the number of overlaps in TF binding. **(G)** Number of regulatory TFs that showed positive expression correlation across protein noise bins. **(H)** Fraction of genes showing SAGA and TFIID dominance across protein noise bins.

This raised a key question - how could an increase in the number of regulatory TFs could lead to increased expression noise. Interestingly, genes regulated by a higher number of TFs showed a concomitant increase in the number of TFs exhibiting cooperative binding (Yang *et al*., 2010; Chen *et al*., 2012) and the genes in the highest noise bins on average had more than 66% cooperative TFs than the genes in the lowest noise bins (Fig. 5C, Supplementary fig. S6C). Expectedly, noise was significantly correlated with the number of cooperatively binding TFs for both mRNA and protein noise (Fig. 5D, Supplementary fig. S6D).

Further, an increase in the number of regulatory TFs and a corresponding increase in their binding sites resulted in a substantial increase in overlap of TF binding sites in the promoter region. The median number of overlaps increased by more than 4-fold for genes in the highest noise bins compared to the genes in the lowest noise bins (Fig. 5E, Supplementary fig. S6E). This was also reflected in the significant correlation between noise at mRNA and protein level with the number TF binding site overlaps (Fig. 5F, Supplementary fig. S6F). Cooperation and competition among TFs can occur only when the TFs are expressed at the same time inside a cell. Interestingly, genes in the highest noise bins on average had more than 90% increase in the number of co-expressing TFs than the genes in the lowest noise bins (Fig. 5G, Supplementary fig. S6G). Further, genes in the highest noise bins had more than 4 times the fraction of SAGA dependent genes and had more than 3 times lower number of TFIID dependent genes compared to the lowest noise bins, considering noise at both mRNA and protein levels (Fig. 5H and Supplementary fig. S6H).

### Cooperative and Competitive TF binding could generate high expression noise

To better understand how an increase in the number of regulatory TFs can lead to higher expression noise, we built a mathematical model of gene regulation and performed stochastic simulations in a population of cells. Specifically, we asked whether a simple increase in the number of regulatory TFs could explain the higher expression noise and whether cooperative and competitive TF binding had any role to play in this process.

We first studied regulation of a gene by a single TF (Fig. 6A). TF binding is a dynamic process consisting of rapid binding and unbinding steps (Burger *et al*., 2010; Das *et al*., 2017). Thus, we used a two-state model of gene expression where a gene could exist in on and off states with specific rates of transition between these two states. The binding of a TF resulted in transition to on-state and thereby led to production of mRNA and proteins. We quantified variations in the gene expression levels over time by stochastic simulations using Gillespie’s algorithm (Fig. 6B). We modeled the dynamics of gene expression in 10000 cells and quantified mean expression level and noise.

**Figure 6.**
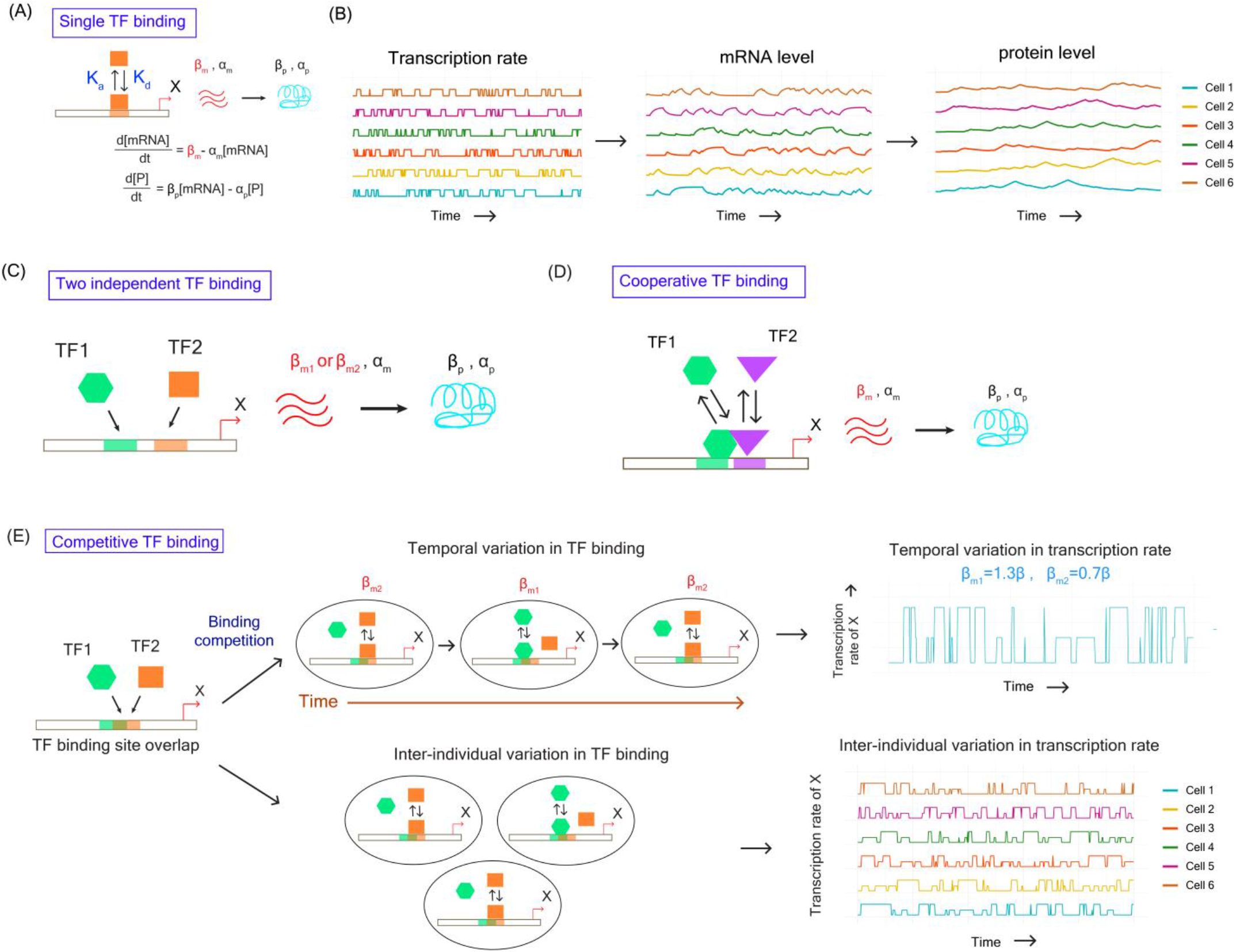
Mathematical modeling and stochastic simulations of TF binding and impact on gene expression noise. **(A)** Mathematical description of the model describing regulation by single TF **(B)** Schematic diagram showing the variation in transcription rate, mRNA levels and protein levels among individual cells obtained from mathematical modeling and stochastic simulation **(C)** Schematic diagram showing gene regulation by two TFs binding independently to the promoter. **(D)** Schematic diagram showing cooperative binding of two TFs to the promoter of a gene and induction of transcription. **(E)** Overlap between TF binding sites lead to binding competition between TFs. This could give rise to temporal variation in TF binding in the same promoter region within a cell. In addition, asynchrony in competitive TF binding among individual cells could give rise to inter-individual variation in TF binding and transcription rate.

In the next step, we tested whether a simple increase in the number of TFs could impact expression noise. To do so, we modeled regulation of a gene by two TFs binding independently to the promoter region (without any cooperation or competition) (Fig. 6C). Here we assumed that binding of any one of the TFs to the promoter led to the on state and resulted in production of mRNA and protein. In case, both the TFs were bound to the promoter, the transcription rate increased and was equal to be the sum of the transcription rates of the individual TFs.

In cooperative TF binding, we modeled the transcription rate by Hill function and assumed that the transcription rate as an all-or-none process regardless of the value of Hill coefficient. This meant that in cooperative binding of two TFs, substantial transcription occurred only when both TFs were simultaneously bound to the promoter region (Fig. 6D). This process could alter the frequency of transcriptional bursts thereby affecting the overall mRNA and protein expression. However, cooperative TF binding can prolong the duration of the on-state and prevent transition to off-state (Gutierrez et al., 2009). We modeled this through a reduction in the off-rate transition. This allowed us to perform all comparisons of expression noise at similar mean expression levels.

Competition among TFs for binding to the overlapping sites in the promoter region could generate noise in two possible ways. First, competition between TFs could lead to a scenario where a gene was regulated by different subsets of TFs at different points of time, thus generating temporal variation (Fig. 6E). In presence of TFs that differ in their strengths of regulation, this could lead to temporal variation in transcription rate in a cell. Secondly, asynchronous temporal variation in TF binding among individual cells in a population could generate inter-individual variation in TF binding (Fig. 6E).

Interestingly, at similar mean protein expression levels, regulation by two independent TFs had lower noise than single TF regulation as the target gene was more frequently in the on state by the action of one of the two TFs and therefore, had less temporal and inter-individual variation in the protein level. This demonstrated that a simple increase in the number of TFs or binding sites could not explain the observed increase in noise (Fig. 7A and 7B). In comparison, both cooperative and competitive binding of TFs led to higher noise compared to regulation by a single TF or by two independent TFs (Fig. 7A and 7B), suggesting that the dynamics of TF binding process in case of gene regulation by multiple TFs had an important role in generation of expression noise.

**Figure 7.**
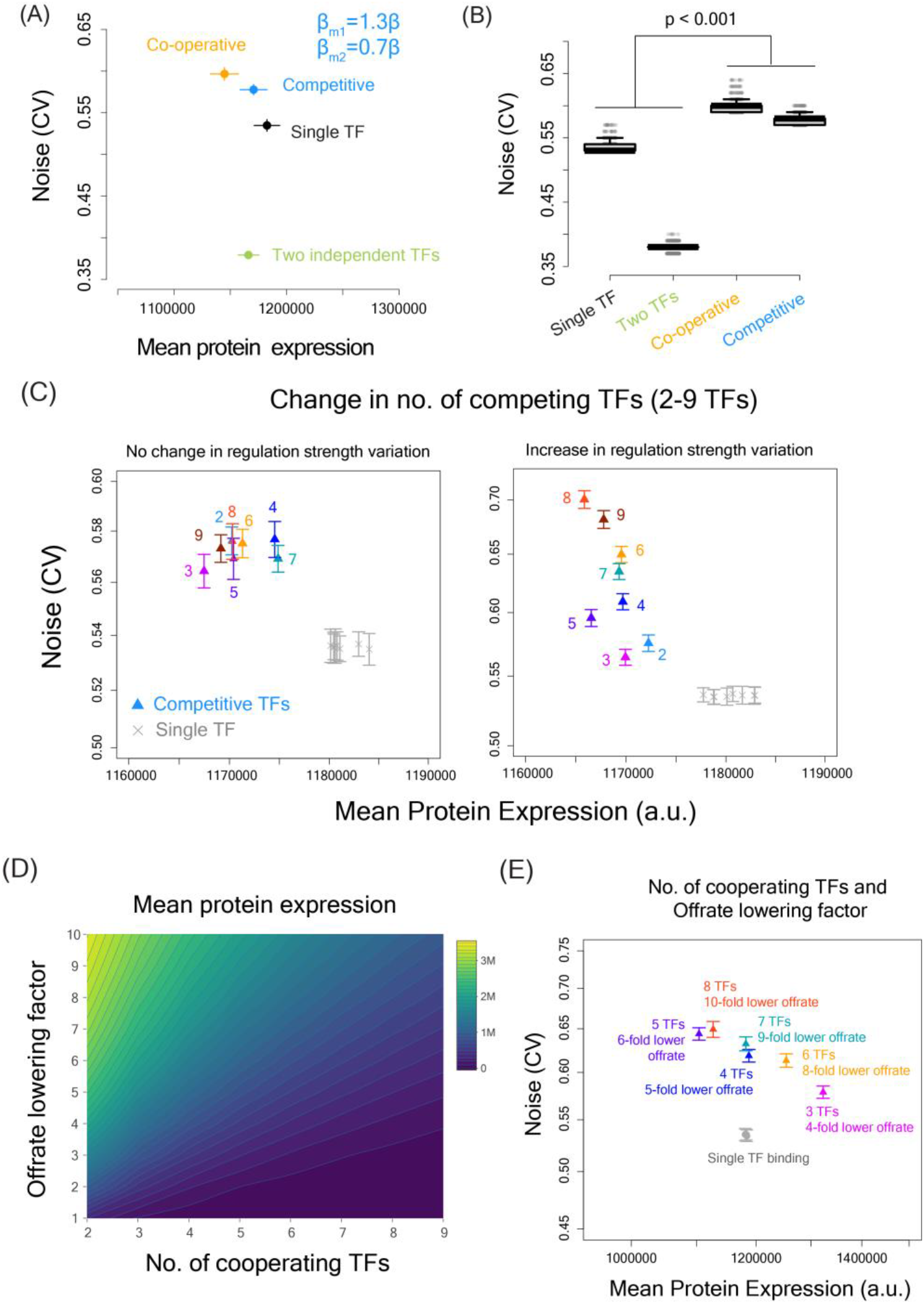
Mean expression and noise in case of gene regulation by a single TF, two independent TFs, two cooperative TFs and two competitive TFs. **(A)** Mean expression and noise values obtained from modeling and simulations of gene regulation by single TF (black), two TFs binding independently (green), two TFs binding cooperatively (orange) and competitively (blue). **(B)** Noise distribution in cases of gene regulation by a single TF and by two TFs binding independently, cooperatively and competitively. Noise was calculated across multiple time-points in 10,000 simulated cells. **(C)** Changes in expression noise with an increase in the number of competitive TFs with or without changes in the variation in regulation strength. **(D)** Increase in the number of cooperating TFs can drive mean protein expression down without any change in the on and off-rates. However, the expression remains the same if binding of more cooperative TFs can increase the time that a gene remains on by lowering the off-rate. **(E)** Gene expression noise in case of 2-9 cooperative TFs at the similar mean protein expression level. We adjusted the off-rate parameter to achieve similar range of mean expression in all cases.

We further explored whether variations in the parameters of the model such as transcription and translation rates, degradation rates, number of cooperative and competitive TFs could influence our inference (Supplementary results). However, over a broad range of parameter values competitive and cooperative TF binding showed higher noise compared to single TF regulation (Supplementary fig. S7-S9).

We extended our analysis to quantify noise in cases of cooperation or competition between more than two TFs as many transcription initiation complexes can contain multiple TFs. Competitive TF binding showed higher noise compared to single TF regulation regardless of the number of TFs and regardless of change in regulation strength among TFs (Fig. 7C). For cooperative TF binding, increase in the number of TFs resulted in a reduction in burst frequency and a reduction in the mean protein level (Supplementary fig. S7). However, an increase in the number of cooperating TFs could proportionally increase the time being spent in the on-state which we modeled through a reduction in off-rate so as to maintain similar mean protein level even with an increase in the number of cooperating TFs (Fig. 7D). In all these scenarios, noise was higher in regulation by cooperative TFs compared to a single TF regulation (Fig. 7E). This suggested that inferences drawn from our models hold regardless of the number of TFs involved in cooperative and competitive binding.

## Discussion

In summary, through an integrated quantitative analysis we have shown that the transcription factor binding process is the most important regulator of gene expression noise. Noisy genes tend to be controlled by a larger number of TFs. These include a substantial fraction of TFs that bind cooperatively to the promoter region. In addition, an increase in the number of regulating TFs can cause an increase in overlap in the TF binding sites which can lead to competition between TFs for binding to the same promoter region. This can give rise to temporal as well as inter-individual variation in TF binding, thereby increasing noise. An earlier work has shown that an increase in the number of transcription factor sites can increase gene expression noise (Sharon *et al*., 2014). However, this study was limited to synthetic promoters and to only a few specific transcription factors. In addition, it was not possible to establish whether the TF binding process was the main driver of noise since the nucleosome occupancy patterns or histone modification patterns for synthetic promoters were unknown. Further, even though competition between specific interacting partners of the TATA binding protein has been implicated in noise before (Ravarani *et al*., 2016), we here describe a much-more general molecular mechanism of noise generation that is not dependent on any specific TF or any specific promoter sequence.

Although our integrated model could predict a substantial fraction of noise variation, there was, however, still a large fraction of noise that could not be explained by our model. This can be due to several reasons. Firstly, it is possible that several other molecular features which can regulate noise have not been considered in our model. Some of these molecular features may still be unknown. Second, there is inherent randomness in molecular processes occurring inside a cell and expression noise can also vary with time. Thus, our calculation of noise at a single time point data may also impact predictive power. Third, the experimental data on molecular features considered in this study have been obtained from different research groups and in different growth conditions. This can impact the predictive ability of our model. Fourth, we understand that some of the features such as the nucleosome occupancy levels and histone modification patterns are dynamic in nature and can change with time. As we modeled these features using datasets obtained at a single timepoint, we might have completely missed the contribution by dynamic nature of these features on noise regulation. Finally, growth conditions, growth rate of cells and cell cycle have all been observed to influence gene expression noise (Zopf *et al*., 2013; Keren *et al*., 2015; Urchueguía *et al*., 2021). Thus, combining data on molecular features across many datasets without consideration for these variables can potentially affect predictive power.

Taken together, our findings provide a step forward for prediction of expression noise. Recent explosion in genomic data has led to genome-wide characterization of TF binding sites across a diverse range of organisms. In addition, with increasing availability of genome-wide nucleosome occupancy maps, histone modification patterns and three-dimensional genome configuration data, our study provides a framework for building integrated model of gene expression noise in other organisms in future. Stochastic variations in molecular processes are ubiquitous in cells across biological systems and have major implications for human diseases. Thus, increased ability to predict variations in biological processes will be extremely useful in quantifying the extent of heterogeneity in cellular traits and phenotypes.

## Methods

### Calculation of expression noise for individual genes

Noise values for individual genes in yeast at the protein level was obtained from Newman *et al*., 2006 and the DM values in the YPD medium were used for all noise analysis. The DM values in the YPD medium were highly correlated with DM values measured in the SD medium and did not affect our results (data not shown). The noise values of all genes at the mRNA level were calculated from the single-cell RNA-seq data provided by Nadal-Ribelles *et al*., 2019 as follows. Briefly, for each gene the coefficient of variation (CV) was calculated from its mean expression and the standard deviation value. Different polynomial fits were made to the CV vs log-transformed mean expression value and the best fit was chosen. A polynomial of order 5 was found to give the best fit. The mean adjusted noise value for a gene was obtained by calculating the vertical distance between the CV value and the best fitted curve. To estimate the impact of outliers on fitting, 95% confidence intervals for the fits were also estimated and were plotted along with the fitted line.

### Building an integrated model of noise in yeast

The integrated model of noise was generated by considering a large number of molecular features that could regulate gene expression. Data on all features were obtained from published experimental datasets. Promoters with the presence of TATA box sequence were identified from the data of Rhee and Pugh, 2012. The genome-wide absolute nucleosome occupancy level for yeast genome was obtained from Oberbeckmann *et al*., 2019 and average nucleosome occupancy level per nucleosome occupied site for all promoter regions and gene bodies were calculated. The region from 1000bp upstream to 10bp downstream of the start codon was considered to be the promoter region of a gene. Average nucleosome occupancy level per occupied site calculated between −1000bp to −900bp region of the start codon was shown as the nucleosome occupancy at −1000bp. Similarly, average nucleosome occupancy level per site was calculated from −900 to −800bp, −800 to −700bp, −700 to −600bp, −600 to −500bp, −500 to −400bp, −400 to −300bp, −300 to −200bp, −200 to −150bp, −150 to −100bp, −100 to −50bp and −50bp to +10bp regions.

Transcription start site (TSS) data for all yeast genes were obtained from Lu and Lin, 2019. Closest TSS site for each gene was obtained, and a spread of potential TSS sites for all genes were calculated. The mRNA synthesis rates and decay rates were obtained from Sun *et al*., 2012. Genome-wide mRNA secondary structure in yeast was obtained from Kertesz *et al*., 2010. The mRNA half-life data and the protein half-life data was obtained from Geisberg *et al*., 2014 and Belle *et al*., 2006 respectively.

Genome-wide histone modification data for yeast was obtained from Pokholok *et al*., 2006 and all modifications were mapped to Gene body, promoter region, and transcription factor binding sites. Histone binding dynamics was obtained from Dion et al., 2007. Data on three-dimensional model of yeast genome was obtained from Duan *et al*., 2010. Number of intra- and Inter-chromosomal contacts for all genes and promoter regions were quantified. tRNA adaptation index for all genes were calculated following the method of Tuller *et al*., 2010. Data on post-translational modifications were obtained from YAAM database (Ledesma *et al*., 2018) and number of different types of modifications for each protein were calculated. For all molecular features, the value of a feature in a biological process was obtained by calculating the median values of the feature for all genes involved in that process.

The classification for SAGA or TFIID dependence of genes for their expression were obtained from Huisinga and Pugh, 2004. In addition, co-activator redundant or TFIID dependent classification of genes were used from Donczew *et al*., 2020. The binding activity of several broadly acting TFs such as TBP, ABF1 and RAP1 were obtained from van Werven *et al*., 2009, Lickwar *et al*., 2012 and de Jonge *et al*., 2020, respectively. Binding patterns of chromatin remodelers were obtained from Yen *et al*., 2012; Zentner and Henikoff, 2013 and Ramachandran *et al*., 2015.

### Analysis of transcription factor binding

The list of transcription factors for all yeast genes were obtained from Yeastract database (http://www.yeastract.com/) (Teixeira *et al*., 2018). For a gene, only those TFs for which experimental evidence for DNA binding had been obtained or the knockout of TF had been experimentally shown to impact expression of the gene were considered. In addition, the binding sites of all TFs to promoter regions of the target genes were searched and mapped using the consensus motif sequences for TFs obtained from YeTFaSCo database (de Boer and Hughes, 2012). All position weighted matrices for all motifs were obtained and all possible combinations of bases were considered. Positions of all such motif sequences of all regulatory TFs of a gene were identified in the promoter region (ranging from −1000bp to +10bp of the start codon) after allowing for maximum two mutations in the consensus sequence. The list of cooperatively binding TFs in yeast were obtained from Yang *et al*., 2010 and Chen *et al*., 2012.

### Regression analysis

The integrated dataset was first scaled using z-score standardization, and the fraction of variation explained and the predictive capability of each molecular feature were quantified by linear regression.

For estimating the predictive power of a class of features on noise, a variable selection method using Ridge and Lasso regression was performed to minimize the problems of multi-collinearity and overfitting. Ridge regression was performed with the R package ‘ridge’ (Cule *et al*., 2020), appropriate number of principal components was chosen and features showing significant effect on noise were identified. Lasso regression was performed using the R package ‘glmnet’ (Friedman *et al*., 2010). The best lambda value was obtained by a 10-fold cross-validation and the lambda for which the cross-validation error was minimum was chosen for subsequent steps. For the preferred lambda value, features whose model coefficients showed non-zero values were considered as features influencing noise. The most important features were further chosen through a stepwise addition and removal of features using stepwise regression where Akaike Information Criterion (AIC) of the fitted models were minimized. This was done using the R package ‘olsrr’ (https://github.com/rsquaredacademy/olsrr). The features in the model with lowest AIC values were selected for further analysis.

In the next step, linear regression with the selected features was performed on the training set to obtain the fraction of variation explained. The linear regression model obtained was then applied on the test data to obtain predicted values along with predicted R^2^ values. The process of dividing dataset, training and prediction was repeated 1000 times to obtain mean and standard deviation values for fraction of variation explained, predicted R^2^ values. Further, random forest models were also built using the selected features using the R package ‘randomForestSRC’ both with and without missing value imputations. These also resulted in fraction of variation explained and predicted R^2^ values.

In addition to the original dataset, two filtered datasets were created with reduced number of variables – one filtered on correlation and another filtered on impact. The first filtered dataset was created by removing features that did not show significant correlation (p<0.05) with noise. To create the second filtered set, first, the impact of all individual features on noise were obtained by linear regression as described above. Only the features that showed significant impact (explained at least 5% of the noise variation or had predicted R^2^ of more than 0.05) were retained in the filtered set. Linear regression was performed on these datasets as described above to obtain fraction of variation explained and R^2^ values for prediction. The analysis showing best results for fraction of variation explained and predicted R^2^ were reported.

### Gene-transcription factor (TF) and TF-TF expression correlation analysis

Gene expression data measured through RNA sequencing from Dhar *et al*., 2019 (NCBI GEO dataset id 104343) was used to calculate all pairwise gene-TF and all pairwise TF-TF (of a gene) expression correlations. Significant positive correlation (p<0.05) between a gene and a regulatory TF indicated that the TF acted as an activator for the gene since the expression of the gene increased with increase in expression of the TF. Similarly, significant negative correlation between a gene and its regulatory TF indicated that the TF acted as a repressor for the gene. The value of correlation coefficient between a gene and a TF (if significant) was taken as the response correlation and the slope of the line was considered as the strength of regulation of the TF. In addition, pairwise expression correlations between all TFs of a gene were calculated. If a TF showed significant positive correlation with at least three other TFs, the TF was considered to be a positively correlated (co-expressing) TF. Similarly, if a TF showed significant negative correlation with at least three other TFs, the TF was considered to be a negatively correlated TF.

### Modeling and stochastic simulation of TF-DNA binding process

The dynamics of TF binding to DNA was studied using a two-state model with consideration for rapid binding and unbinding of TF to DNA. The binding-unbinding of TF to DNA was considered to be a Poisson process and thus, the time intervals between two successive bindings (or two successive unbindings) were exponentially distributed. The time intervals between successive events (on or off switching) were sampled from exponential distributions with rate parameters λ_on_ and λ_off_ respectively. The dynamics of cooperative binding and competitive binding of TFs was compared to the dynamics of regulation by a single TF and two independent TFs. For modeling binding of two TFs on- and off-time intervals were sampled from Poisson distributions individually for each of the TFs with same rate parameters. For cooperative binding, only when both the TFs were bound to the promoter, the gene switched to the on state and led to production to mRNA and protein molecules at the same rates as the single TF binding. This resulted in lowering of burst frequency in case of cooperative TF binding which eventually reduced the mean protein level. To address this issue, λ_off_ was gradually reduced to achieve similar mean expression level as in single TF regulation. Reduction of λ_off_ values prolong the on-state cooperative TF binding (Gutierrez *et al*., 2009). Competitive TF binding was modeled in the same way as modeling single TF binding but TF that bound to the promoter at every on-state transition was chosen. The rate of production of mRNA was influenced by the regulation strength of the TF that bound to the promoter.

Binding of a TF led to switching to on state which resulted in production of mRNA at a rate β_m_ and translation of these mRNA molecules to proteins at a rate β_p_. These mRNA and protein molecules were considered to undergo removal resulting from dilution due to cell growth and degradation at the rates of α_m_ and α_p_ respectively.

The dynamics of transcription and translation were modeled using the following equations.

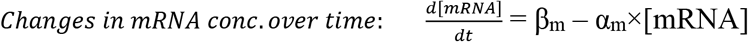

where β_m_ denoted the transcription rate per unit time (or burst size) and αm denoted the removal rate of mRNA due to degradation and dilution.

Similarly,

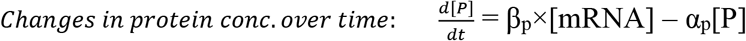

where β_p_ denoted the protein production rate from mRNA and α_p_ denoted the protein removal rate.

The parameters β_p_, α_m_, α_p_ were considered to be same across all cases of TF binding. In case of cooperative TF binding, β_m,coop_ was assumed to be the same as the β_m,single_. In case of competitive TF binding, the production rates varied between two TFs, with β_m1_ = l.3×β_m,single_ and β_m2_ = O.7×β_m,single_.

All rate parameters for single TF, two independent TF, cooperative TF and competitive TF binding were chosen in such a way that the comparisons were mathematically equivalent. As the concentrations of TFs can impact the chances of binding, the concentration of TF in single TF binding scenario was considered to be same as the concentration of each of the cooperatively binding TFs. Further, the concentration of the TF in single TF binding scenario was considered to be equal to the sum of the concentrations of two TFs in case of competitive binding scenario. The transcription rates in the cases of regulation by single TF and by cooperative TFs were same. The transcription rates in case of regulation by two independent TFs were chosen in such a way that the average transcription rate was same as the transcription rate in regulation by a single TF. The transcription rates of the TFs in case of competitive TFs were chosen in such a way that the average transcription rate of the two TFs was same as the transcription rate in single TF regulation.

Stochastic simulations were performed using Gillespie’s algorithm (Gillespie, 1977) to decipher the dynamics of TF-DNA binding in all three scenarios and to investigate the impact of cooperative and competitive TF binding on noise. The behavior of the system was tracked at small discrete time intervals Δt from the initial time point t. These resulted in observations at n time points t, t+ Δt, t+2×Δt, …, t+n×Δt. Any event of binding or unbinding occurring within a time interval was noted and resulted in changes in transcription rate which eventually led to a change in protein concentration (Supplementary fig. 5B). Binding of TFs led to transcription and increase in mRNA and protein concentration according to the above equations. As the time interval Δt was considered to be small, the equations modeling the behavior of the systems was simplified as

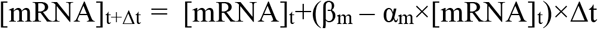

and

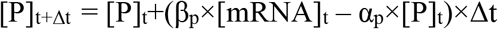

β_m_, α_m_, β_p_ and α_p_ were expressed in the units of Δt for further simplification of these equations. The dynamics of transcription, variation in the mRNA concentration and variation in protein concentration with time were modeled across 10,000 cells (Supplementary fig. S5B). Noise was expressed as coefficient of variation (CV) from the calculation of mean and standard deviation in the protein level across these 10,000 cells and across all individual time points.

Mean expression level and noise values were calculated for a wide range of parameter values for all the parameters λ_on_, λ_off_, β_m_, β_p_, α_m_, and α_p_ to ensure that the results obtained were not biased by the choice of specific parameter values. All noise comparisons were made at similar mean expression levels to eliminate any bias in the noise values due to variations in mean expression levels.

## Supporting information

Complete supplementary information

## Availability of data and materials

The datasets analysed during the current study are available in the NCBI-GEO repository. Complete supplementary information is available at FigShare.

Code availability - https://github.com/riddhimandhar/IntegratedNoiseModel

## Authors’ contributions

RD conceived the study, RD, LP, and SP performed data analysis, RD wrote the manuscript with inputs from LP and SP. All authors read and approved the manuscript.

## Acknowledgments

We are extremely grateful to Dr. Ben Lehner for his insightful and critical comments on the first draft of the manuscript.

