## Supplementary material for "Transcription factor binding process is the primary driver of noise in gene expression": Complete supplementary information

This PDF file contains

**Supplementary Results**

**Supplementary figures S1 to S12**

**Supplementary table S1**

### Supplementary Results

#### Cooperative and Competitive TF binding caused higher noise across a wide range of model parameter values

As genes could vary in their switching rates between on and off states, mRNA and protein synthesis rates and removal rates, we further explored how changes in these parameters would impact mean expression level and noise. We first quantified how changes in switching rates impacted the mean expression level in cases of single TF, cooperative TF and competitive TF regulations (Supplementary fig. S7A). This assumed significance as we aimed to perform all comparisons of noise values among these three regulation scenarios at similar mean expression levels. Changes in on and off -rates resulted in alterations in burst frequency (Supplementary fig. S7B, C). Some of the combinations of on- and off-rates resulted in very low or very high burst frequency and led to situations where genes were mostly off or mostly on respectively (Supplementary fig. S7B, C). These resulted in very low noise irrespective of the gene regulation mechanism. Thus, to avoid these scenarios, we carefully chose the on- and off-rate parameters so as to remain within a reasonable burst frequency range. Variations in mRNA and protein synthesis rates and degradation rates altered the mean expression levels but across all parameter ranges noise values were substantially higher in the cases of competitive and cooperative TF binding compared to single TF regulation (Supplementary fig. S8, S9A). Expectedly, in case of competitive TF binding, increase in variation in regulation strength of competing TFs led to further increase in noise (Supplementary fig. S8). Changes in transcription on and off-rates also altered noise levels, but again for competitive TF binding, noise was higher compared to single TF regulation (Supplementary fig. S8). For cooperatively binding TFs, changing on and off-rates resulted in substantial divergence in mean protein level compared to single TF regulation. Therefore, we compared the noise levels across all on- and off-rates and observed higher noise in case of cooperative TF binding (Supplementary fig. S9B).

Overlaps in binding sites for different TFs can also lead to degeneracy of TF binding sequences to accommodate different consensus binding motifs of different TFs. Such degeneracy can change binding affinity of TFs to the DNA and can lead to noisy transcription. However, we did not see any difference in binding site degeneracy among high and low- noise genes (Supplementary fig. S10).

**Noise analysis in biological processes**

Further, even though noise in expression of individual genes has been well-studied, the architecture of noise in cellular pathways has remained largely unexplored beyond the traditional gene-set enrichment analysis. A majority of the cellular traits and phenotypes are polygenic in nature, meaning they are shaped by coordinated functioning of groups of genes. Fluctuations in the expression of such group of genes shape the variations in the activity of cellular pathways, thereby generating phenotypic heterogeneity. Thus, it is essential to decipher the pattern of noise at the pathway levels to quantitatively understand variations in cellular phenotypes.

To do so, we associated genes to biological processes according to the Gene ontology classification for *S. cerevisiae* obtained from http://geneontology.org/ (data version: releases/2017-11-22) (Ashburner *et al.*, 2000; Carbon *et al.*, 2019) and mapped gene names to GO terms using SGD yeast gene association map (https://www.yeastgenome.org/, GOC Validation Date: 11/10/2017) (Cherry *et al.*, 2012). We considered only terms associated with Biological Processes (BP) in our analysis and a process was included in our analysis only if gene expression noise data for at least 10 genes were available for that class. For each included GO term, we measured noise dispersion by calculating inter-quartile range of noise values of genes participating in that process (Supplementary fig. S11A). Noise value for each individual gene was obtained from mean adjusted noise value calculated from single-cell RNA-seq data (Nadal-Ribelles *et al.*,

2019) as well as from distance to median (DM) measure calculated for protein noise data (Newman *et al., 2*006). We were able to obtain noise dispersion values for 475 biological processes based on mRNA noise data and for 192 processes based on protein noise data. To identify processes with high or low noise, we performed one-sample Wilcoxon signed-rank test to check whether the noise values of genes in a biological process differed significantly from zero followed by multiple hypothesis testing correction using Benjamini-Hochberg method (Benjamini and Hochberg, 1995) with a False Discovery Rate (FDR) cut-off of 10%.

Biological processes showed a wide range of distributions for noise dispersion values (Supplementary fig. S11B,C). Interestingly, majority of the biological processes (>50%), however, did not have any significant deviation towards high or low noise and were balanced in their content of high- and low-noise genes (Wilcoxon rank-sum test, FDR<0.1). Processes showing high noise both based on mRNA noise data and protein noise data included cellular pathway associated with oxidative stress response. Common processes that exhibited low noise in both mRNA and protein noise datasets were associated with transcription and translation.

**Regulatory TF binding process also predicted noise in biological processes**

Noise in biological processes - shaped by the noise levels of individual genes - has important implications for phenotypic heterogeneity. Thus, we investigated how each of the features used in the analysis of expression noise at individual gene level could explain the variation in the data and could predict noise across biological processes. We did so for both mRNA and protein noise data and ranked them. Interestingly, all of the top 10 features for explaining and predicting noise at the gene level could also explain and predict noise of the biological processes to a large extent (Supplementary fig. S12A). Most interestingly, the number of regulatory TFs and the number of TF binding sites were among the top predictors of noise dispersion across biological processes (Supplementary fig. S12A).

Again, TF binding process was the single most important contributor to the faction of variation explained and predictive power of all the features combined. The integrated model comprising of all features was able to explain 53-69% the noise dispersion of the biological processes and was able to predict noise dispersion with predicted $R^2$ value of 0.4-0.57 (Supplementary fig. S12B). As before, TF binding alone could explain 49-58% of the variation in noise dispersion across biological processes and was able to predict noise dispersion with predicted $R^2$ value of 0.32-0.44 (Supplementary fig. S12B). These results highlighted a pivotal role of the regulatory TFs in predicting noise across biological processes.

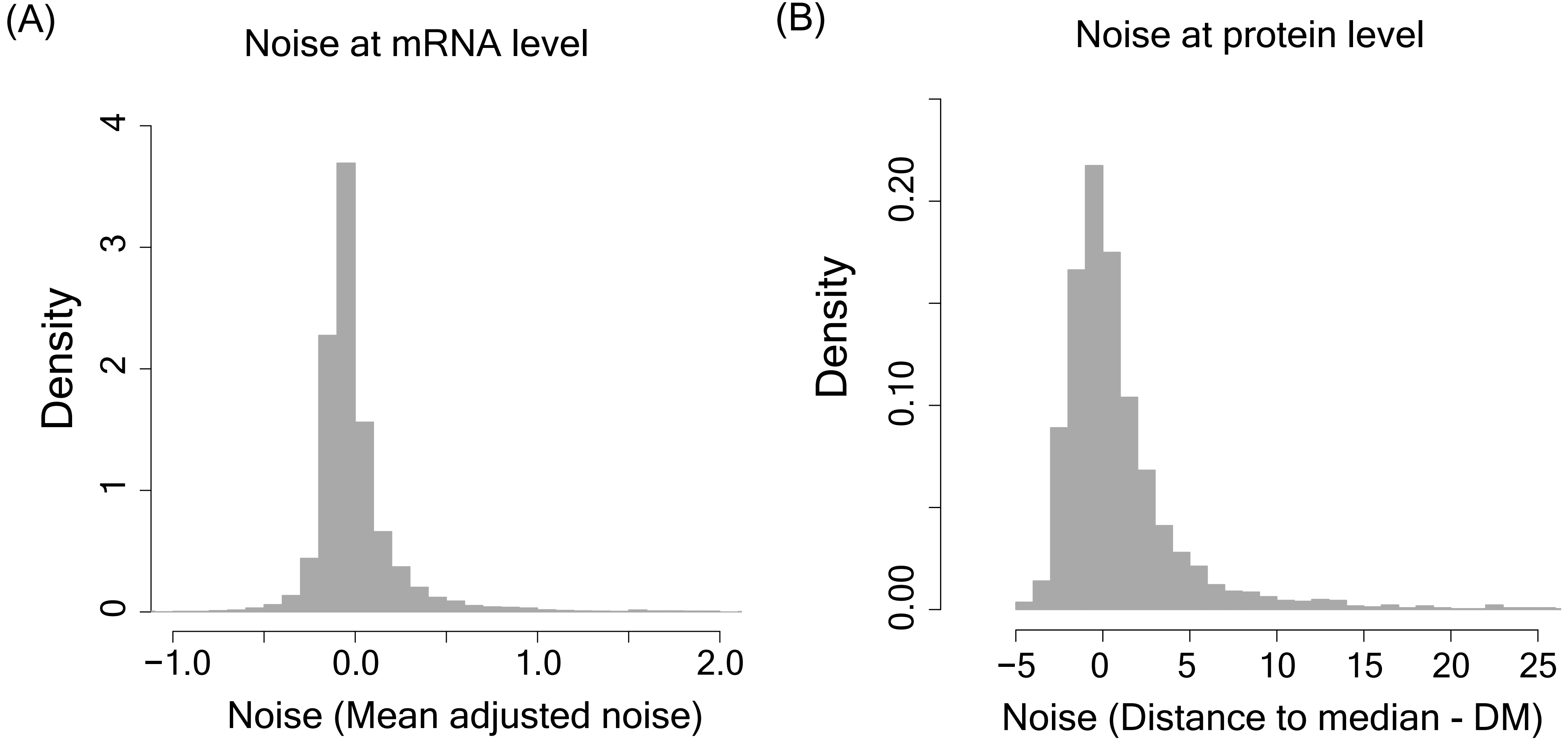


**Supplementary figure S1:** Distribution of expression noise of all genes at the mRNA level **(A)** and at the protein level **(B)**.

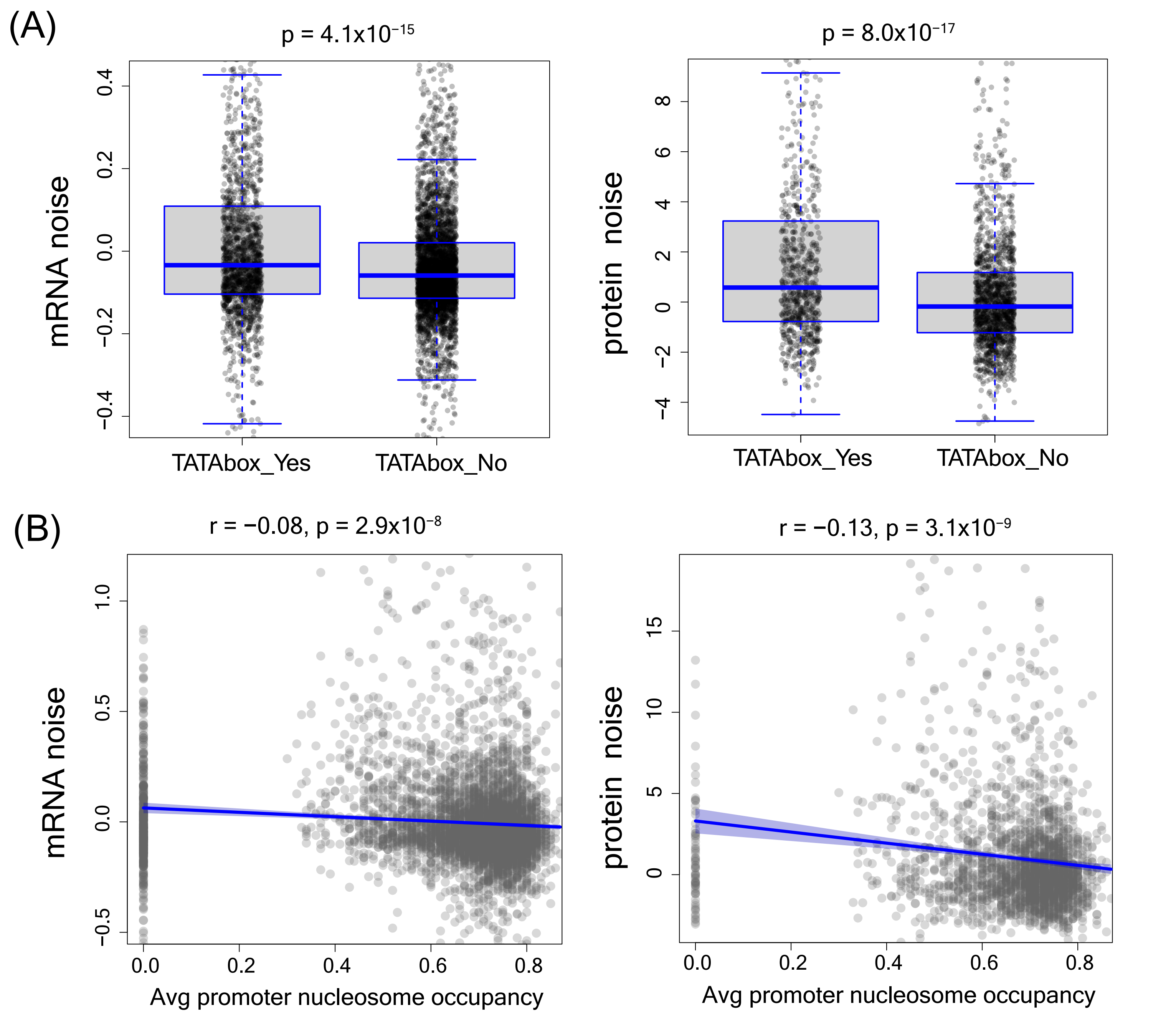


**Supplementary figure S2: (A)** Difference in expression noise of genes with and without the TATA box sequence in the promoter, calulated at the mRNA as well as the protein level **(B)** Correlation between noise and average promoter nucleosome occupancy.

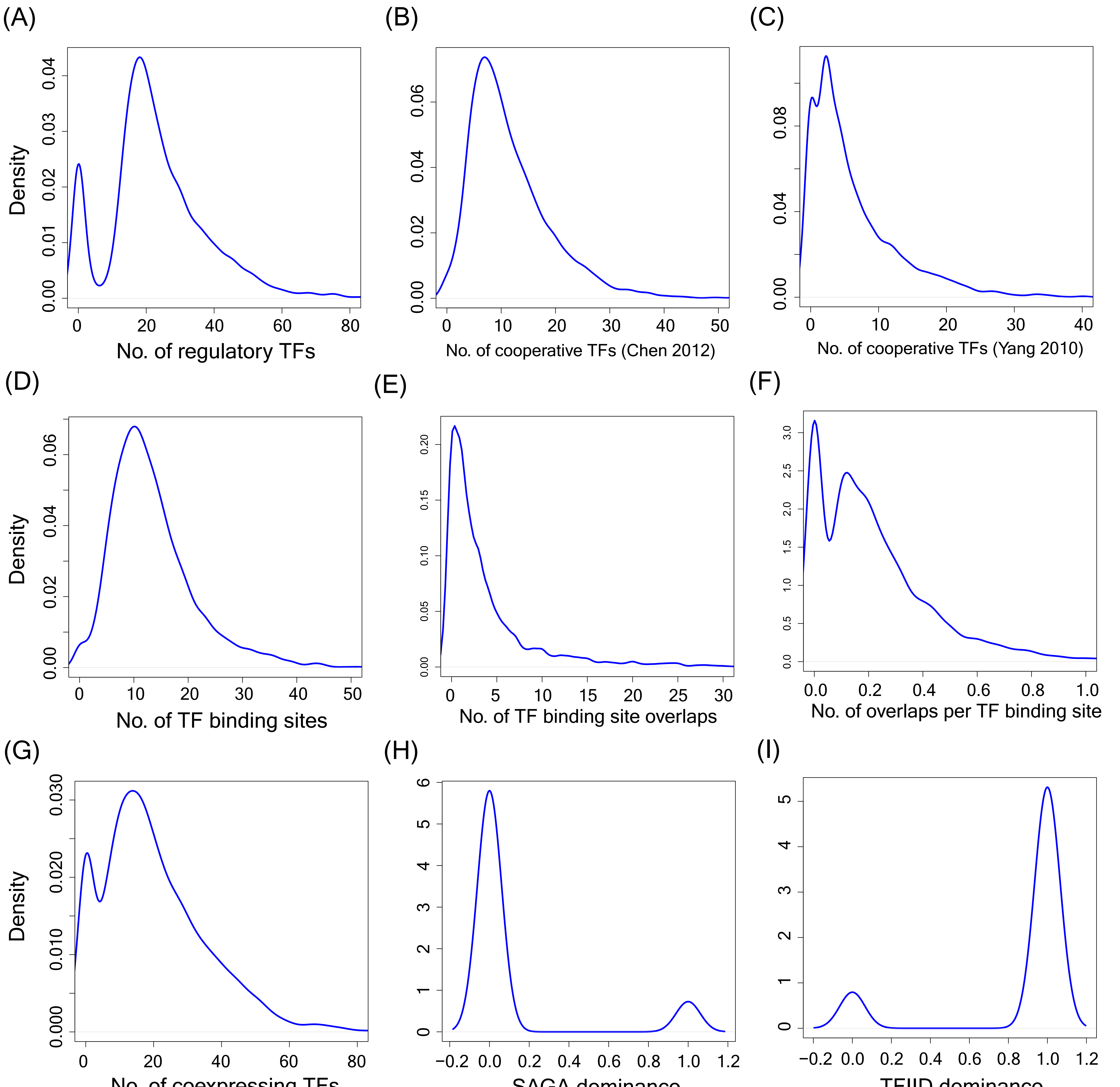


**Supplementary figure S3 - Distributions of feature values**. Plots showing distributions of **(A)** number of reguatory TFs, **(B-C)** number of cooperative TFs (Chen et al., 2012 and Yang et al., 2010), **(D)** number of TF binding sites, **(E)** number of TF binding site overlaps, **(F)** number of overlaps per TF binding site, **(G)** number of TFs showing positive expression correlation among themselves, **(H)** number of genes with SAGA dominance in the promoter, and **(I)** number of genes with TFIID dominance in the promoter.

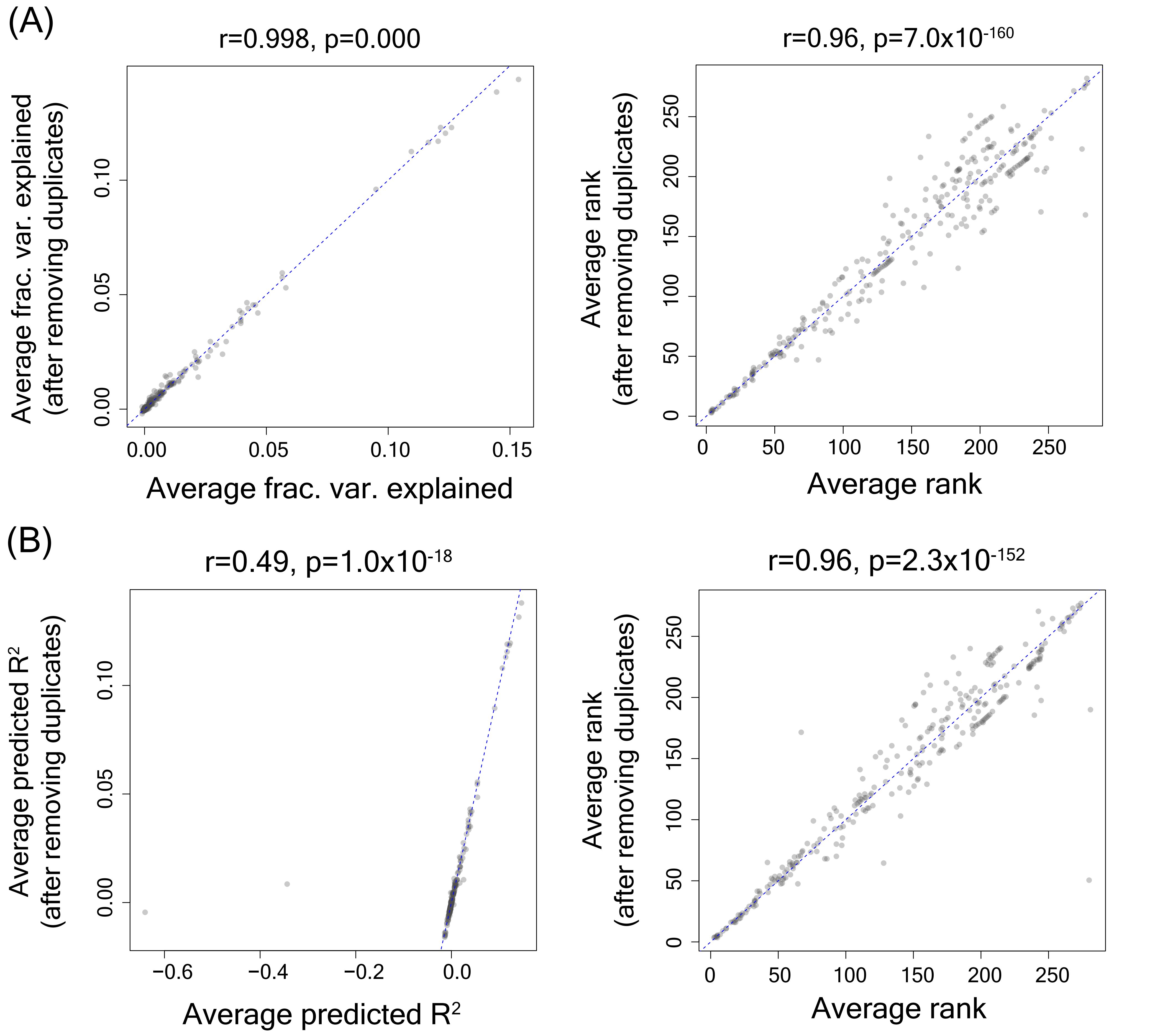


**Supplementary figure S4 - Comparison of variation explained and predictive ability of the model between datasets with and without duplicate genes. (A)** Correlation between average fraction variation explained and the average rank for all features with and without removing duplicate genes from the data. Average fraction variation explained and average rank for the same were calculated by calculating average values from mRNA and protein noise data. **(B)** Correlation between average predicted $R^2$ and the corresponding average rank with and without removing duplicate genes from the data.

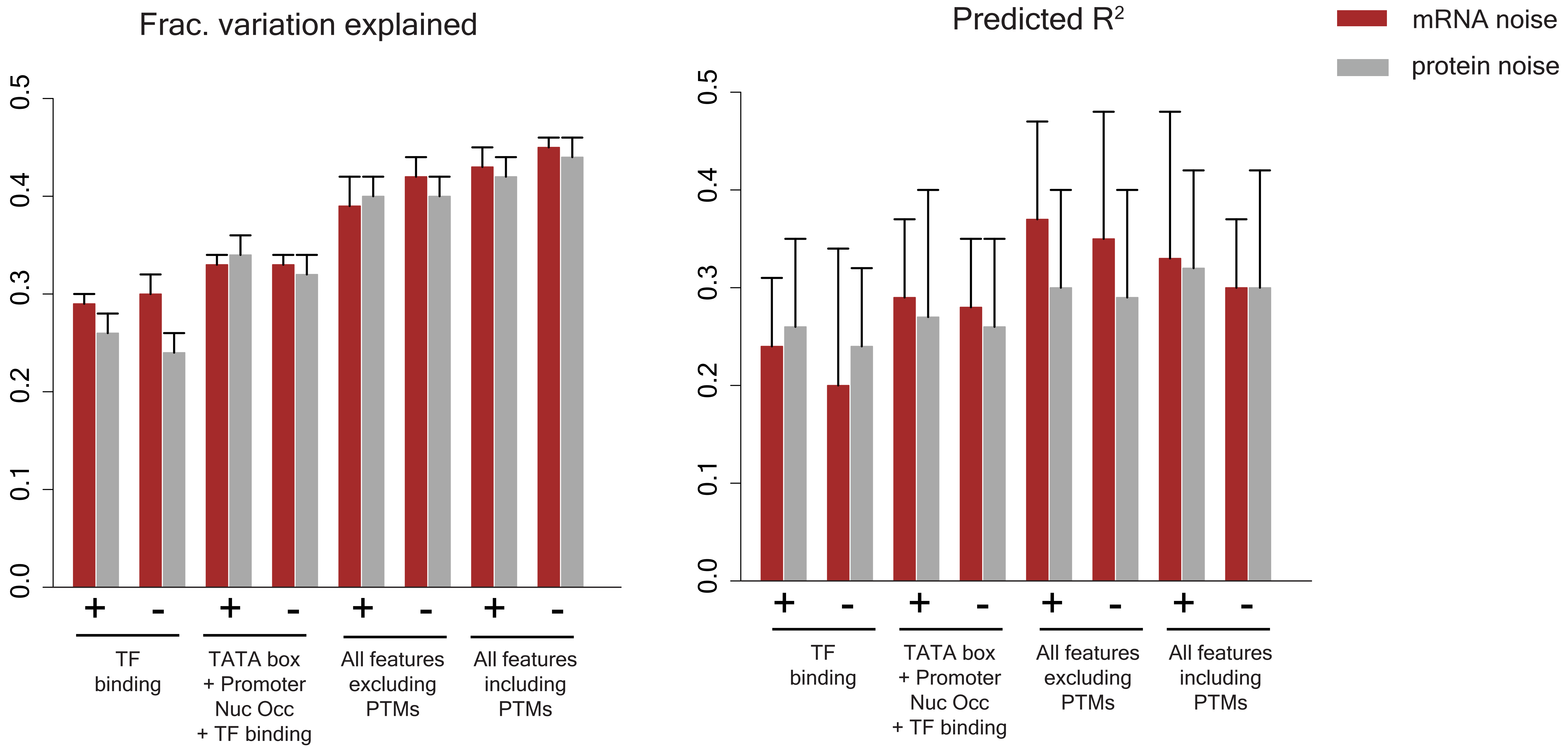


**Supplementary figure S5** - Fraction of variation explained in noise data and predictive ability (given by predicted $R^2$ value) by features associated with TF binding activity, combination of TF binding activity with other features, combination of all features excluding PTMs and combination of all features including PTMs. The '+' and '-' signs denote the datasets used in analysis, with '+' indicating the full dataset and the '-' sign indicating the dataset after removal of duplicate genes.

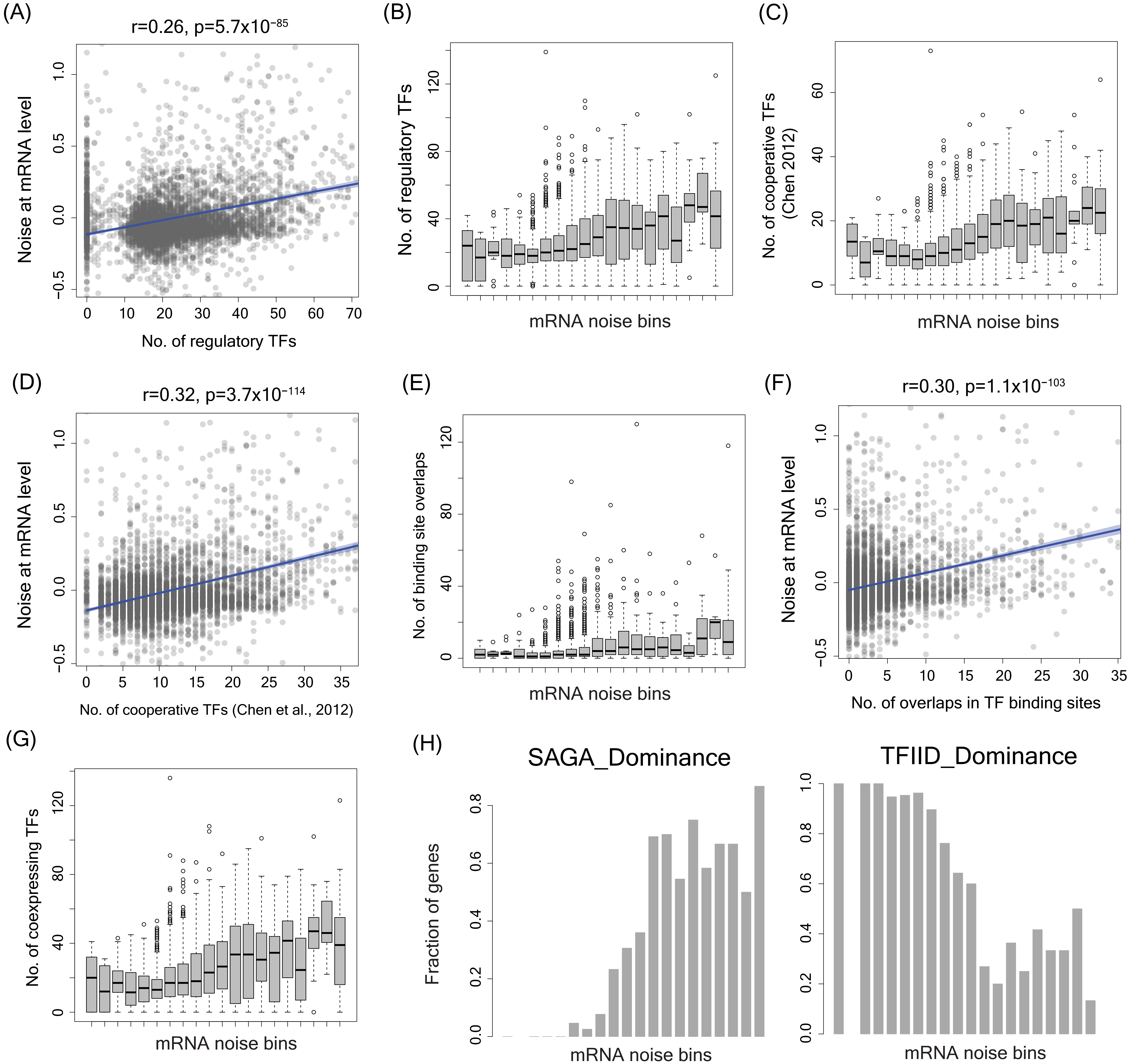


**Supplementary fig. S6 - Genes with high noise were regulated by a higher number of TFs, had higher number of cooperatively binding TFs, and showed more overlaps in TF binding sites compared to low-noise genes. (A)** Correlation between noise at mRNA level and the number of regulatory TFs **(B)** Number of regulatory TFs of genes across different mRNA noise bins. **(C)** Number of cooperative TFs of genes across mRNA noise bins. **(D)** Correlation between noise at mRNA level and the number of cooperative TFs (Chen et al., 2012) **(E)** Number of overlaps between TF binding sites for genes across mRNA noise bins. **(F)** Correlation between noise at mRNA level and the number of overlaps in TF binding sites **(G)** Number of regulatory TFs that showed positive expression correlation across mRNA noise bins. **(H)** Fraction of genes showing SAGA and TFIID dominance across mRNA noise bins.

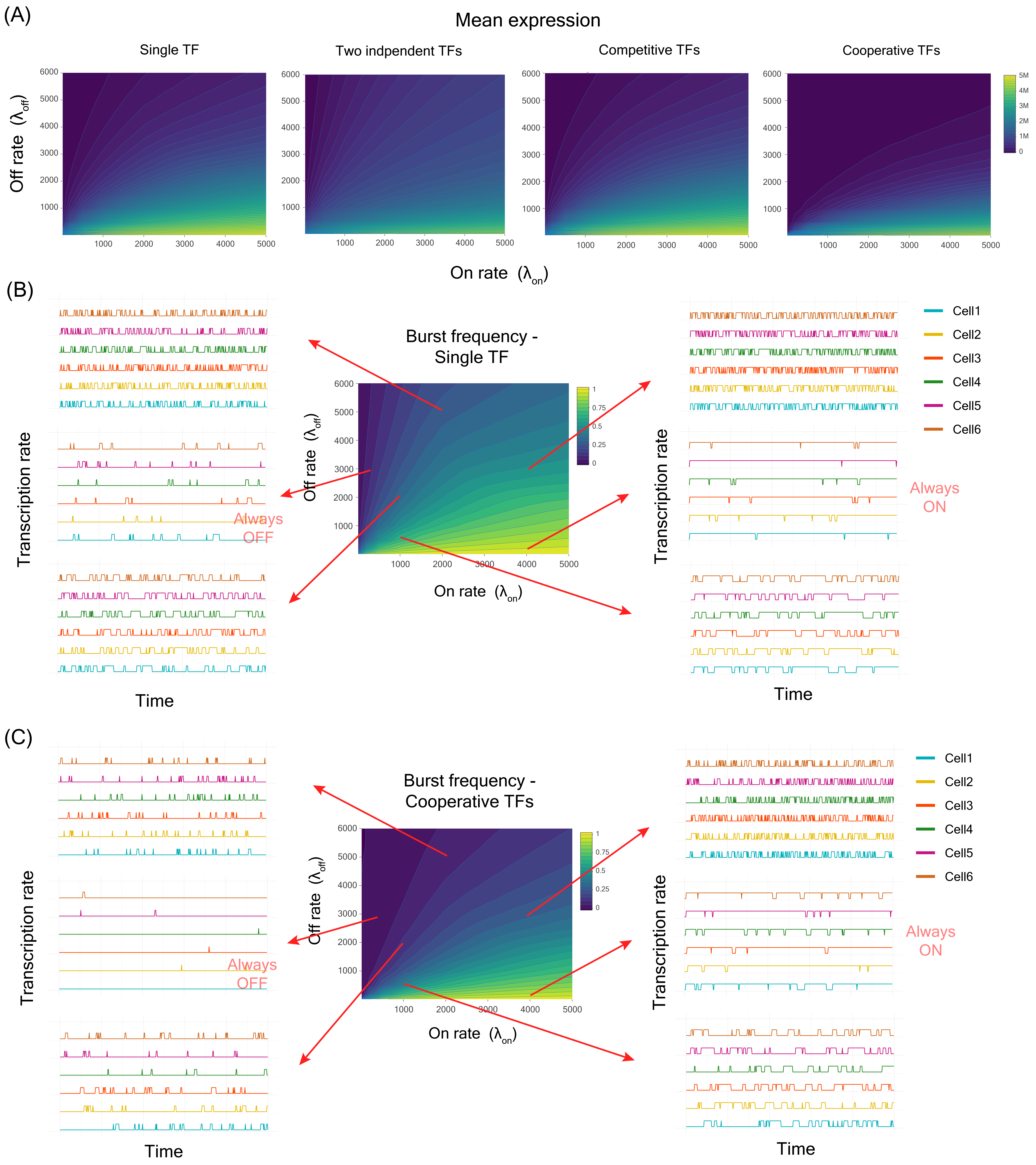


**Supplementary figure S7**: **Changes in on- and off-rate parameters impact burst frequency and influence mean expression level.** **(A)** Relationship between on- and off-rate parameters ($\lambda_{on}$ and $\lambda_{off}$ respectively)and mean expression levels in cases of regulation by single TF, two independent TFs, competitive TFs and cooperative TFs. **(B)** Variation in transcription rate over time (burst frequency) in single TF regulation **(C)** Variation in transcription rate over time (burst frequency) in case of regulation by cooperatively binding TFs. For same values of $\lambda_{on}$ and $\lambda_{off}$, the cells are in always on or always off states.

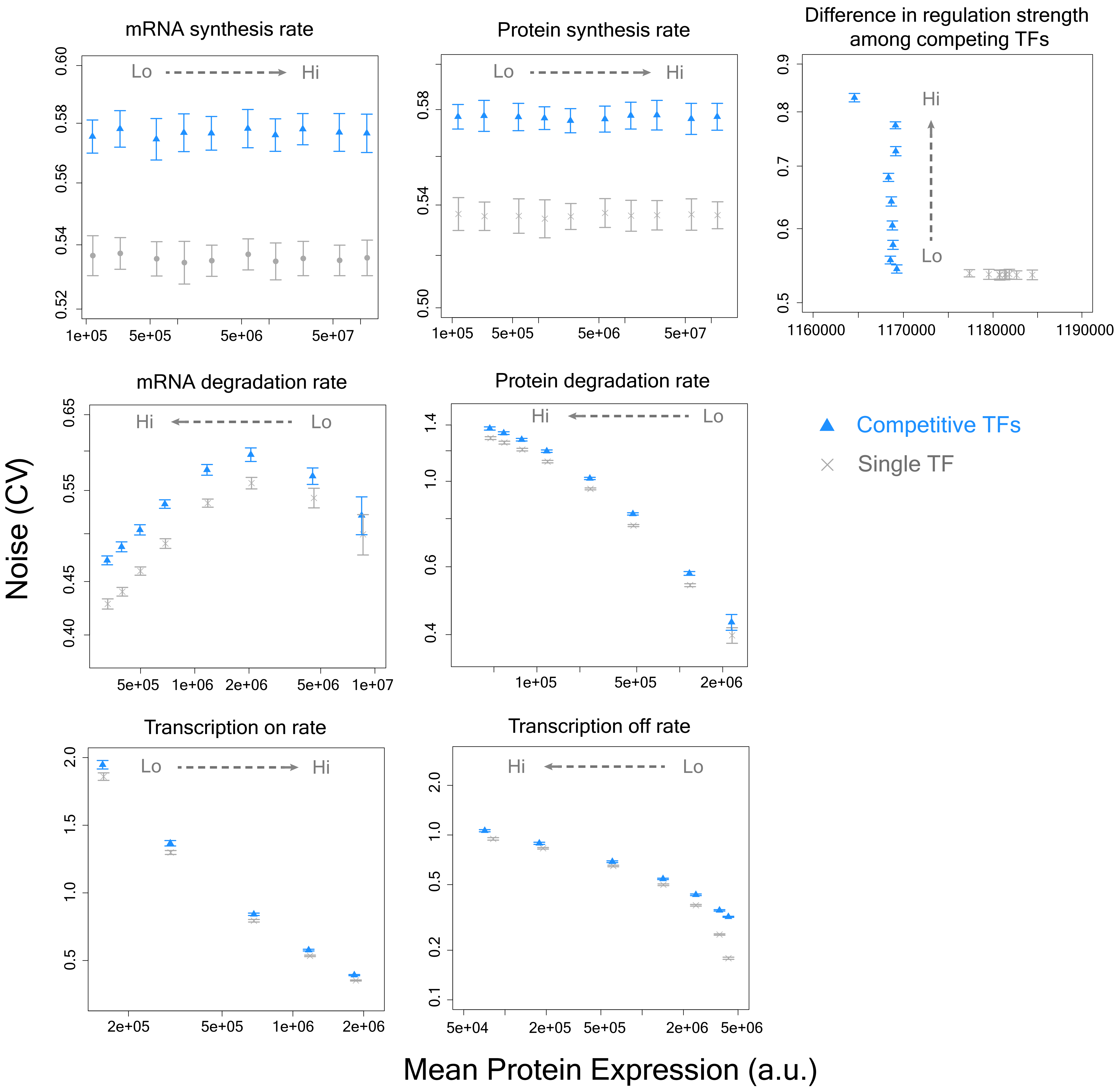


**Supplementary figure S8: Noise in case of competitive TF binding is higher compared to single TF regulation across a wide range of parameter values.** Changes in mRNA and protein synthesis rates, mRNA and protein degradation rates, on- and off-rate parameters ($\lambda_{on}$ and $\lambda_{off}$ respectively) changes mean expression levels both in single TF and competitive TF binding, but the noise levels in competitive binding are always higher than single TF binding. Inceased variation in regulatory strengths of competitive TFs lead to higher noise.

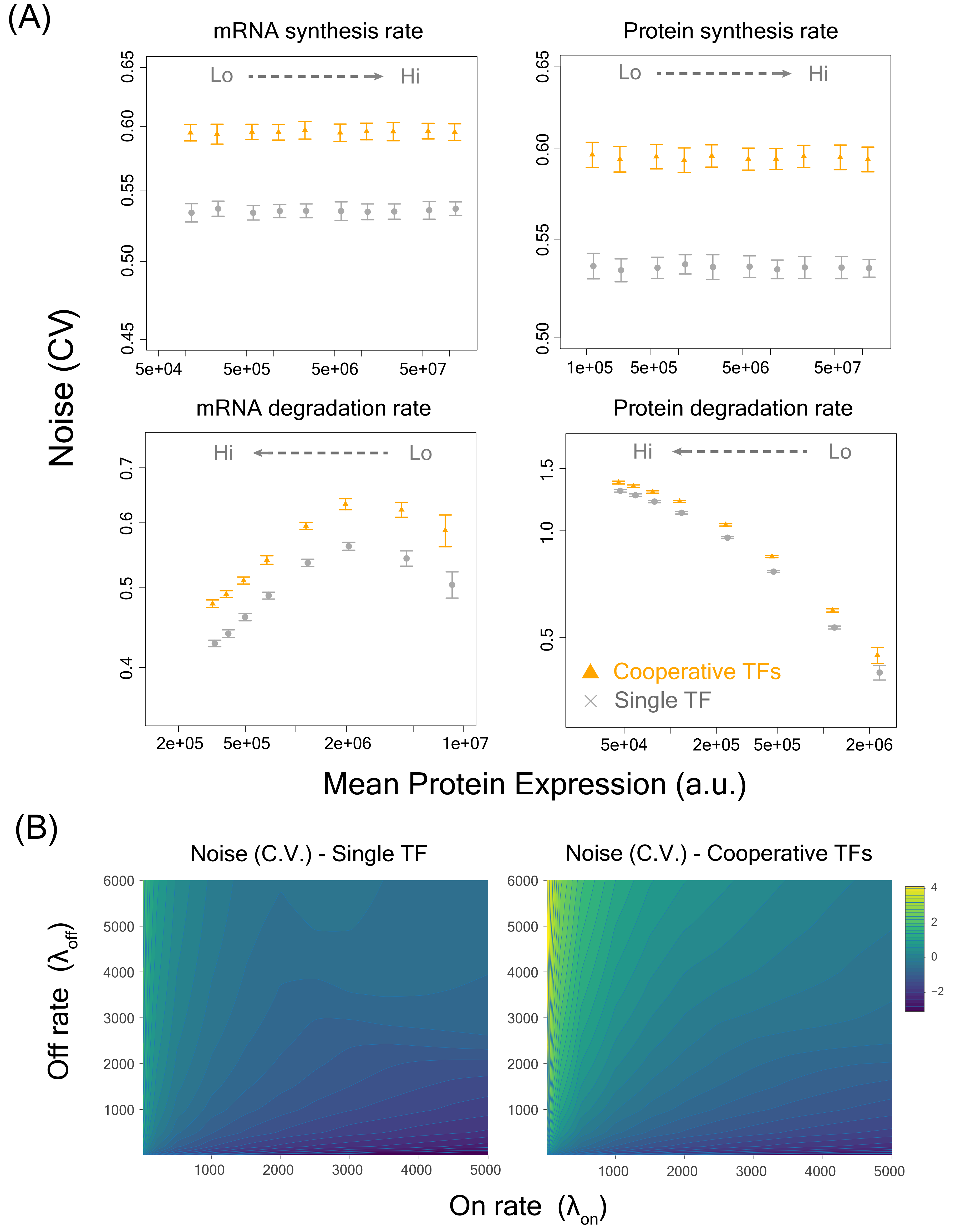


**Supplementary figure S9**: **Noise in case of cooperatively binding TFs is higher compared to single TF regulation across a wide range of parameter values. (A)** Changes in mRNA and protein synthesis rates, mRNA and protein degradation rates changes mean expression levels both in single TF and cooperative TF binding, but the noise levels in cooperative binding are always higher than single TF binding. **(B)** Noise values across a wide range of on- and off-rate parameter values ($\lambda_{on}$ and $\lambda_{off}$ respectively) for single TF and cooperative TF binding.

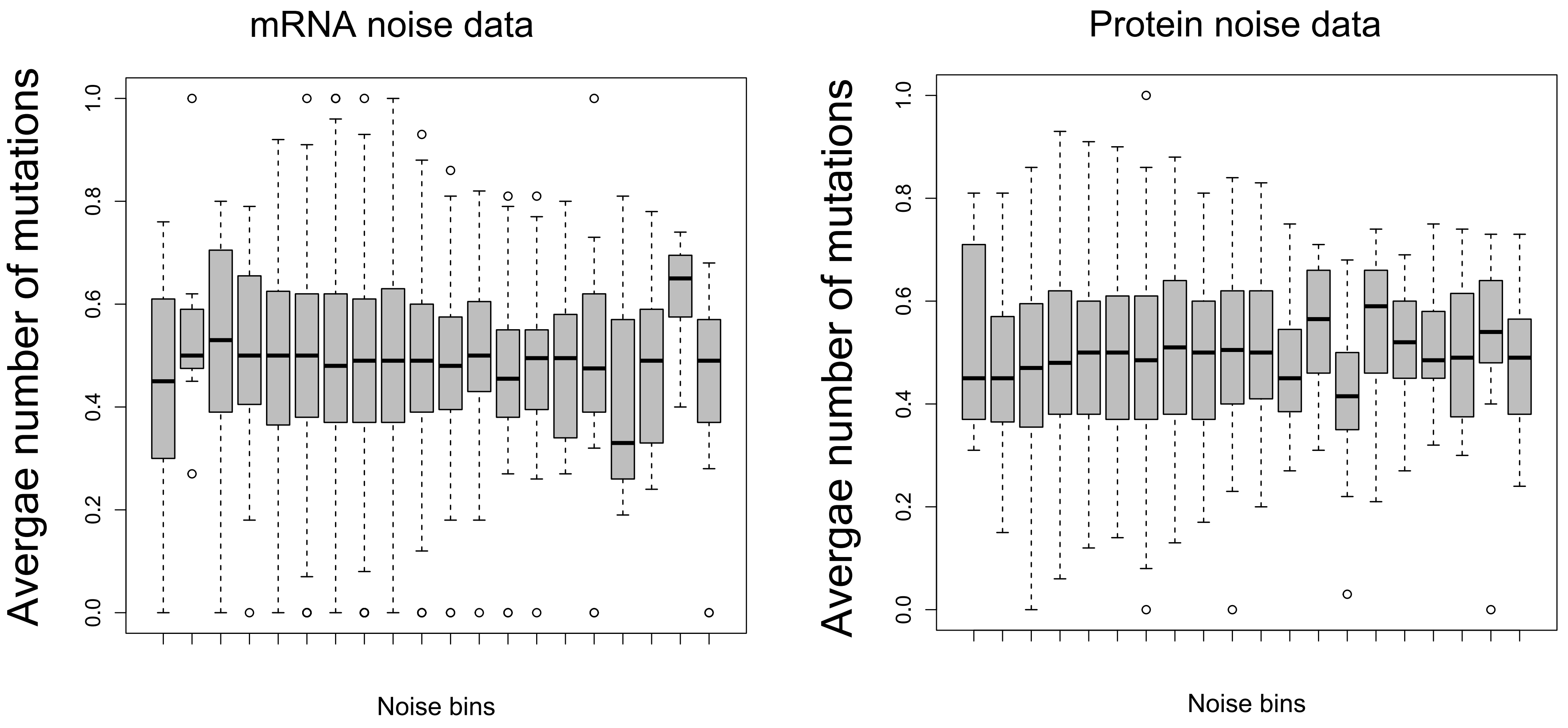


**Supplementary figure S10**: No difference in the average number of mutations in the overlapping TF binding sites between low- and high-noise genes, based on calculations both at the mRNA and protein levels.

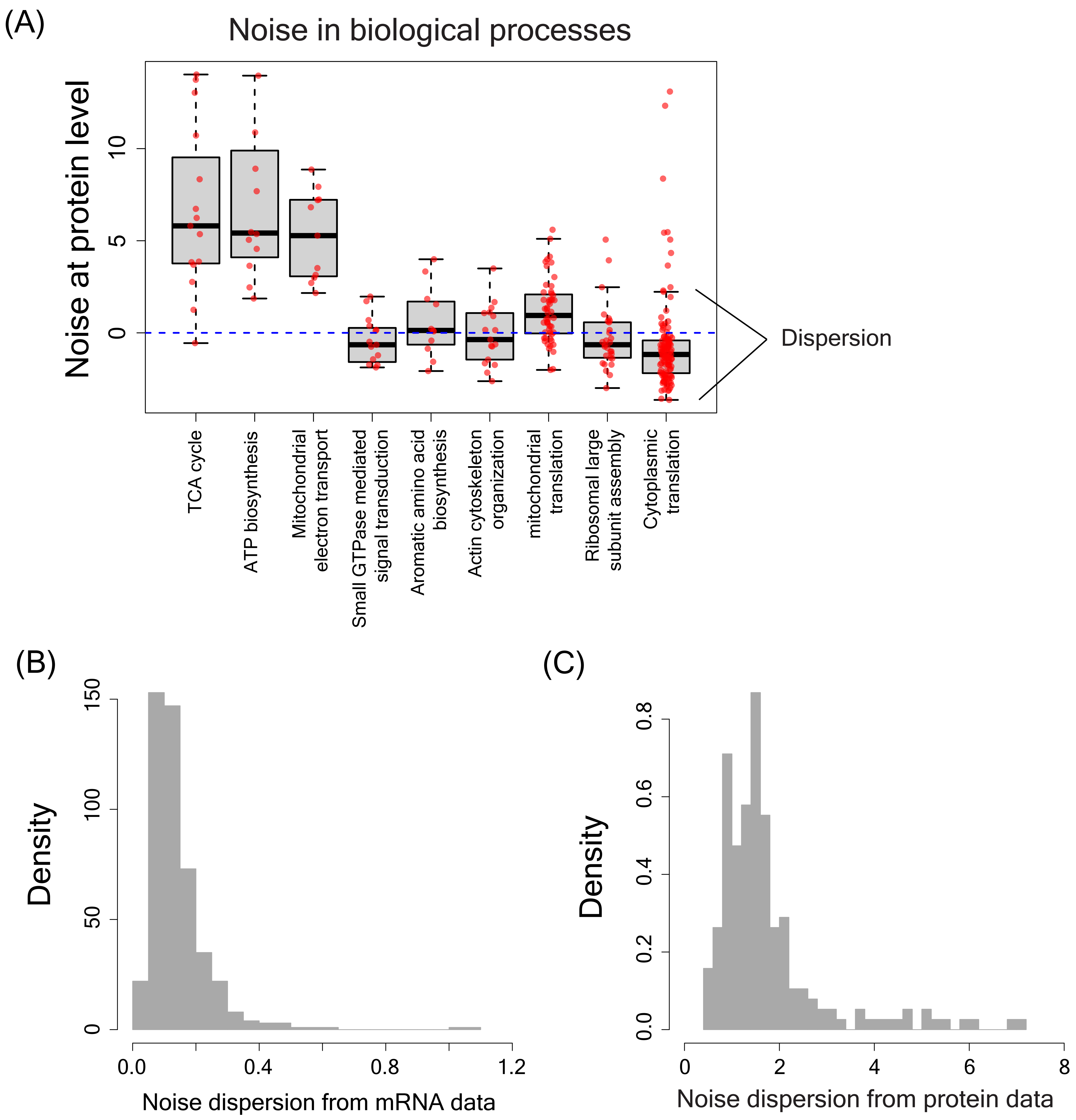


**Supplementary figure S11: (A)** Examples of noise distribution in several biological processes with noise calculated at the protein level. Distribution of noise dispersion values of biological processes calculated from noise data at the mRNA level **(B)** and from noise data at the protein level **(C)**

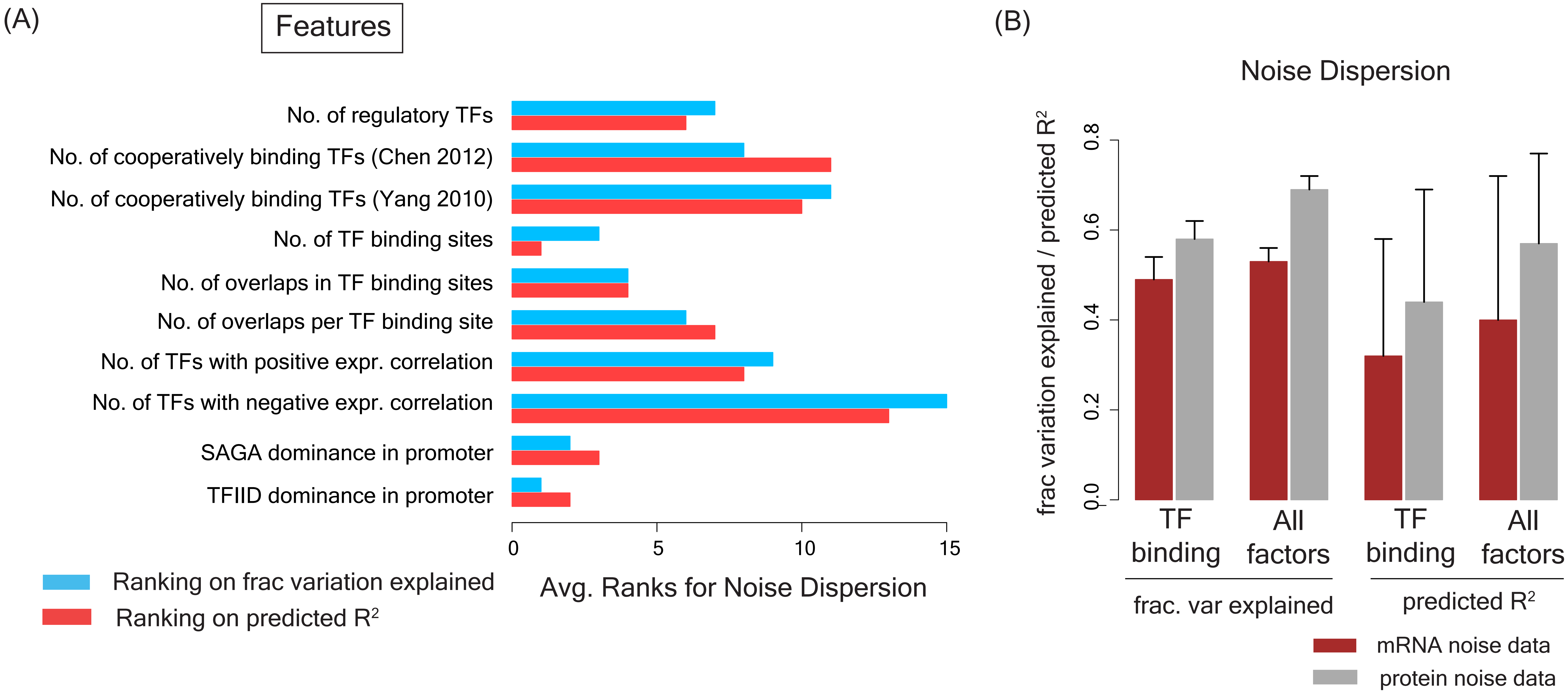


**Supplementary figure S12** - TF binding activity also predicts noise across biological processes **(A)** Average ranking of features based on the fraction of variation explained and by predictive ability (given by the predicted $R^2$ value) for noise dispersion across biological processes, calculated based on both mRNA and protein noise. **(B)** Fraction of variation explained in noise across biological processes and predictive ability by features associated with TF binding activity, and by the combination of all features.

### Supplementary Table S1: List of features included in our integrated model of noise

| Feature name | Description | References |
|---|---|---|
| STRE_elem | Presence/Absence of the Stress Response Element in the promoter | Moskvina et al., Yeast 1998 |
| TATAbox | presence/Absence of the TATAbox sequence in the promoter | Rhee and Pugh, Nature 2012 |
| NumPromNucOcc | Number of sites in the promoter occupied by nucleosomes | Oberbeckmann et al., Genome Res. 2019 |
| AvPromNucOcc | Average absolute nucleosome occupancy level per nucleosome bound site in the promoter | |
| LenAvgPromNucOcc | Average length of nucleosome occupancy in the promoter | |
| NumGeneNucOcc | Number of sites in the genebody occupied by nucleosomes | |
| AvGeneNucOcc | Average absolute nucleosome occupancy level per nucleosome bound site in the genebody | |
| N_PrNO50 | Number of sites upto 50 bp upstream of the start codon occupied by nucleosomes | |
| N_PrNO100 | Number of sites between 50 bp and 100bp upstream of the start codon occupied by nucleosomes | |
| N_PrNO150 | Number of sites between 100 bp and 150bp upstream of the start codon occupied by nucleosomes | |
| N_PrNO200 | Number of sites between 150 bp and 200bp upstream of the start codon occupied by nucleosomes | |
| N_PrNO300 | Number of sites between 200 bp and 300bp upstream of the start codon occupied by nucleosomes | |
| N_PrNO400 | Number of sites between 300 bp and 400bp upstream of the start codon occupied by nucleosomes | |
| N_PrNO500 | Number of sites between 400 bp and 500bp upstream of the start codon occupied by nucleosomes | |
| N_PrNO600 | Number of sites between 500 bp and 600bp upstream of the start codon occupied by nucleosomes | |
| N_PrNO700 | Number of sites between 600 bp and 700bp upstream of the start codon occupied by nucleosomes | |
| N_PrNO800 | Number of sites between 700 bp and 800bp upstream of the start codon occupied by nucleosomes | |
| N_PrNO900 | Number of sites between 800 bp and 900bp upstream of the start codon occupied by nucleosomes | |
| N_PrNO1000 | Number of sites between 900 bp and 1000bp upstream of the start codon occupied by nucleosomes | |
| V_PrNO50 | Level of absolute nucleosome occupancy in the region upto 50bp upstream of the start codon | |
| V_PrNO100 | Level of absolute nucleosome occupancy in the region between 50bp and 100bp upstream of the start codon | |
| V_PrNO150 | Level of absolute nucleosome occupancy in the region between 100bp and 150bp upstream of the start codon | |
| V_PrNO200 | Level of absolute nucleosome occupancy in the region between 150bp and 200bp upstream of the start codon | |
| V_PrNO300 | Level of absolute nucleosome occupancy in the region between 200bp and 300bp upstream of the start codon | |
| V_PrNO400 | Level of absolute nucleosome occupancy in the region between 300bp and 400bp upstream of the start codon | |
| V_PrNO500 | Level of absolute nucleosome occupancy in the region between 400bp and 500bp upstream of the start codon | |
| V_PrNO600 | Level of absolute nucleosome occupancy in the region between 500bp and 600bp upstream of the start codon | |
| V_PrNO700 | Level of absolute nucleosome occupancy in the region between 600bp and 700bp upstream of the start codon | |
| V_PrNO800 | Level of absolute nucleosome occupancy in the region between 700bp and 800bp upstream of the start codon | |

| | | |
|---|---|---|
| V_PrNO900 | Level of absolute nucleosome occupancy in the region between 800bp and 900bp upstream of the start codon | |
| V_PrNO1000 | Level of absolute nucleosome occupancy in the region between 900bp and 1000bp upstream of the start codon | |
| whetherTF | Whether the gene is a TF (Yes/No) | Yeastract database; YeTFaSCo database; Pokholok et al., Cell 2005; Newman et al., Nature 2006; Dhar et al., eLife, 2019 |
| Num_RegTF_YeastractYT | Number of regulatory TFs (from Yeastract data) | |
| MedExp_TFYT | Median expression of regulatory TFs (Yeastract data) | |
| MedDM_SD_TFYT | Median noise of regulatory TFs (Yeastract data) | |
| MedPosDM_SD_TFYT | Median positive noise (DM values) of regulatory TFs (Yeastract data) | |
| MedNegDM_SD_TFYT | Median negative noise (DM values) of regulatory TFs (Yeastract data) | |
| PercNegDM_TFYT | Percentage of TFs showing negative noise (DM) values | |
| PercPosDM_TFYT | Percentage of TFs showing positive noise (DM) values | |
| minDM_TFYT | Minimum noise (DM) value | |
| maxDM_TFYT | Maximum noise (DM) value | |
| PosCorTF_YT | Number of TFs showing positive expression correlation with the target gene | |
| NegCorTF_YT | Number of TFs showing negative expression correlation with the target gene | |
| PosCorTF.NegCorTF_YT | Ratio of the number of TFs showing positive expression correlation to the number of TFs showing negative expression correlation with the target gene | |
| PercPosCorTF_YT | Percentage of TFs showing positive expression correlation with the target gene | |
| PercNegCorTF_YT | Percentage of TFs showing negative expression correlation with the target gene | |
| Both_PosCorr_NegCorrTF_YT | Whether the gene has positively and negatively correlated TFs (Yes/No) | |
| NoisePosCorTF_YT | Noise (DM) value of TFs showing positive expression correlation with the target gene | |
| NoiseNegCorTF_YT | Noise (DM) value of TFs showing negative expression correlation with the target gene | |
| MeanStrPosCorTF | Mean regulation strength of positively correlated TFs | |
| SdStrPosCorTF | Sd regulation strength of positively correlated TFs | |
| MeanStrNegCorTF | Mean regulation strength of negatively correlated TFs | |
| SdStrNegCorTF | Sd regulation strength of negatively correlated TFs | |
| MeanCorPosCorTF | Mean correlation value of TFs showing positive expression correlation with the target gene | |
| SdCorPosCorTF | Sd correlation value of TFs showing positive expression correlation with the target gene | |
| MeanCorNegCorTF | Mean correlation value of TFs showing negative expression correlation with the target gene | |
| SdCorNegCorTF | Sd correlation value of TFs showing negative expression correlation with the target gene | |
| NumPosCor_withinTFs | Number of TFs showing positive expression correlation with other TFs regulating the same target gene | |
| NumNegCor_withinTFs | Number of TFs showing negative expression correlation with other TFs regulating the same target gene | |
| PercPosCor_withinTFs | Percentage of TFs showing positive expression correlation with other TFs regulating the same target gene | |
| PercNegCor_withinTFs | Percentage of TFs showing negative expression correlation with other TFs regulating the same target gene | |

| | |
|---|---|
| PosCor.NegCor_withinTFs | Ratio of the number of TFs showing positive expression correlation with other regulating TFs of a gene to the number of TFs showing negative expression correlation with other TFs of a gene |
| PercPosCorWN_OVsites | Percentage of TFs showing positive expression correlation with other TFs regulating the same target gene and binding to overlapping binding sites in the promoter |
| PercNegCorWN_OVsites | Percentage of TFs showing negative expression correlation with other TFs regulating the same target gene and binding to overlapping binding sites in the promoter |
| AvgMut | Average number of mutations in the TF binding motifs in the promoter region |
| N_TFSites100 | Number of TF binding sites upto 100bp upstream region of the start codon |
| N_TFSites200 | Number of TF binding sites within 100bp and 200bp upstream region of the start codon |
| N_TFSites300 | Number of TF binding sites within 200bp and 300bp upstream region of the start codon |
| N_TFSites400 | Number of TF binding sites within 300bp and 400bp upstream region of the start codon |
| N_TFSites500 | Number of TF binding sites within 400bp and 500bp upstream region of the start codon |
| N_TFSites600 | Number of TF binding sites within 500bp and 600bp upstream region of the start codon |
| N_TFSites700 | Number of TF binding sites within 600bp and 700bp upstream region of the start codon |
| N_TFSites800 | Number of TF binding sites within 700bp and 800bp upstream region of the start codon |
| N_TFSites900 | Number of TF binding sites within 800bp and 900bp upstream region of the start codon |
| N_TFSites1000 | Number of TF binding sites within 900bp and 1000bp upstream region of the start codon |
| ExpTF100 | Mean expression of TFs binding upto 100bp upstream region of the start codon |
| NoiseTF100 | Expression noise of TFs binding upto 100bp upstream region of the start codon |
| ExpTF200 | Mean expression of TFs binding between 100bp and 200bp upstream region of the start codon |
| NoiseTF200 | Expression noise of TFs binding between 100bp and 200bp upstream region of the start codon |
| ExpTF300 | Mean expression of TFs binding between 200bp and 300bp upstream region of the start codon |
| NoiseTF300 | Expression noise of TFs binding between 200bp and 300bp upstream region of the start codon |
| ExpTF400 | Mean expression of TFs binding between 300bp and 400bp upstream region of the start codon |
| NoiseTF400 | Expression noise of TFs binding between 300bp and 400bp upstream region of the start codon |
| ExpTF500 | Mean expression of TFs binding between 400bp and 500bp upstream region of the start codon |
| NoiseTF500 | Expression noise of TFs binding between 400bp and 500bp upstream region of the start codon |
| ExpTF600 | Mean expression of TFs binding between 500bp and 600bp upstream region of the start codon |
| NoiseTF600 | Expression noise of TFs binding between 500bp and 600bp upstream region of the start codon |
| ExpTF700 | Mean expression of TFs binding between 600bp and 700bp upstream region of the start codon |

| | |
|---|---|
| NoiseTF700 | Expression noise of TFs binding between 600bp and 700bp upstream region of the start codon |
| ExpTF800 | Mean expression of TFs binding between 700bp and 800bp upstream region of the start codon |
| NoiseTF800 | Expression noise of TFs binding between 700bp and 800bp upstream region of the start codon |
| ExpTF900 | Mean expression of TFs binding between 800bp and 900bp upstream region of the start codon |
| NoiseTF900 | Expression noise of TFs binding between 800bp and 900bp upstream region of the start codon |
| ExpTF1000 | Mean expression of TFs binding between 900bp and 1000bp upstream region of the start codon |
| NoiseTF1000 | Expression noise of TFs binding between 900bp and 1000bp upstream region of the start codon |
| PercTFsiteNuc | Percentage of TF sites showing nucelosome occupancy |
| PercTFsiteHistMod | Percentage of TF sites with histone modifications |
| AvgTFsiteNucOcc | Average TF site nucleosome occupancy level |
| AvgTFsiteHist | Average TF site histone level |
| AvgTFsiteMod | Average TF site histone modifications |
| AvgTFsiteAsoc | Average level of associated regulators and modifiers (GCN4,GCN5,ESA1) in TF binding sites |
| PercOfPromNuc | Percentage of the total promoter nucleosome occupancy level observed in the TF binding sites |
| PercOfPromHist | Percentage of the total promoter histone level observed in the TF binding sites |
| PercOfPromMod | Percentage of the total promoter histone modifications observed in the TF binding sites |
| PercOfPromAsoc | Percentage of the total promoter associated regulators and modifiers observed in the TF binding sites |
| TFsiteH3 | H3 level in TF binding sites |
| TFsiteH4 | H4 level in TF binding sites |
| TFsiteH3K9ac_vsH3 | Level of H3K9ac modifications in TF binding sites |
| TFsiteH3K14ac_vsH3 | Level of H3K14ac modifications in TF binding sites |
| TFsiteH4ac_vsH3 | Level of H4ac modifications in TF binding sites |
| TFsiteH3K4me1_vsH3 | Level of H3K4me1 modifications in TF binding sites |
| TFsiteH3K4me2_vsH3 | Level of H3K4me2 modifications in TF binding sites |
| TFsiteH3K4me3_vsH3 | Level of H3K4me3 modifications in TF binding sites |
| TFsiteH3K36me3_vsH3 | Level of H3K36me3 modifications in TF binding sites |
| TFsiteH3K79me3_vsH3 | Level of H3K79me3 modifications in TF binding sites |
| TFsiteESA1 | Level of ESA1 in TF binding sites |
| TFsiteGCN5 | Level of GCN5 in TF binding sites |
| TFsiteGCN4.AA | Level of GCN4 in TF binding sites |
| PercOfPromH3 | Percentage of the total promoter H3 level observed in the TF binding sites |
| PercOfPromH4 | Percentage of the total promoter H4 level observed in the TF binding sites |
| PercOfPromH3K9ac_vsH3 | Percentage of the total promoter H3K9ac level observed in the TF binding sites |
| PercOfPromH3K14ac_vsH3 | Percentage of the total promoter H3K14ac level observed in the TF binding sites |
| PercOfPromH4ac_vsH3 | Percentage of the total promoter H4ac level observed in the TF binding sites |
| PercOfPromH3K4me1_vsH3 | Percentage of the total promoter H3K4me1 level observed in the TF binding sites |
| PercOfPromH3K4me2_vsH3 | Percentage of the total promoter H3K4me2 level observed in the TF binding sites |
| PercOfPromH3K4me3_vsH3 | Percentage of the total promoter H3K4me3 level observed in the TF binding sites |

| | |
|---|---|
| PercOfPromH3K36me3_vsH3 | Percentage of the total promoter H3K36me3 level observed in the TF binding sites |
| PercOfPromH3K79me3_vsH3 | Percentage of the total promoter H3K79me3 level observed in the TF binding sites |
| PercOfPromESA1 | Percentage of the total promoter ESA1 level observed in the TF binding sites |
| PercOfPromGCN5 | Percentage of the total promoter GCN5 level observed in the TF binding sites |
| PercOfPromGCN4.AA | Percentage of the total promoter GCN4 level observed in the TF binding sites |
| NumSites | Number of TF binding sites in the promoter region |
| NumOverlaps | Number of overlaps in TF binding sites in the promoter region |
| RatOverlap.Numsites | Ratio of the number of overlaps to the total number of TF binding sites |
| AvgOverlapLen | Average overlap length |
| AvgfrOverlapLen | Average fraction of TF binding site showing overlap |
| OV100 | Percentage of binding site overlaps up to 100bp upstream region of the start codon of all overlaps in the promoter |
| OV200 | Percentage of binding site overlaps between 100bp and 200bp upstream region of the start codon |
| OV300 | Percentage of binding site overlaps between 200bp and 300bp upstream region of the start codon |
| OV400 | Percentage of binding site overlaps between 300bp and 400bp upstream region of the start codon |
| OV500 | Percentage of binding site overlaps between 400bp and 500bp upstream region of the start codon |
| OV600 | Percentage of binding site overlaps between 500bp and 600bp upstream region of the start codon |
| OV700 | Percentage of binding site overlaps between 600bp and 700bp upstream region of the start codon |
| OV800 | Percentage of binding site overlaps between 700bp and 800bp upstream region of the start codon |
| OV900 | Percentage of binding site overlaps between 800bp and 900bp upstream region of the start codon |
| OV1000 | Percentage of binding site overlaps between 900bp and 1000bp upstream region of the start codon |
| PercAct2_Overlap | Percentage of overlapping sites shared by two activators |
| Avg_stract2_ov | Average strength of regulation of two activators binding to overlapping sites |
| Df_stract2_ov | Difference in strength of regulation of two activators binding to overlapping sites |
| Avg_corstr_act2_ov | Average expression correlation of two activators binding to overlapping sites with the target gene |
| Df_corstr_act2_ov | Difference in expression correlation of two activators binding to overlapping sites with the target gene |
| PercRep2_overlap | Percentage of overlapping sites shared by two repressors |
| Avg_strrep2_ov | Average strength of regulation of two repressors binding to overlapping sites |
| Df_strrep2_ov | Difference in strength of regulation of two repressors binding to overlapping sites |
| Avg_corstr_rep2_ov | Average expression correlation of two repressors binding to overlapping sites with the target gene |
| Df_corstr_rep2_ov | Difference in expression correlation of two repressors binding to overlapping sites with the target gene |
| PercActrep_overlap | Percentage of overlapping sites shared by one activator and one repressor |
| Avg_stractrep_ov | Average strength of regulation of activator and repressor binding to overlapping sites |

| | | |
|---|---|---|
| Df_stractrep_ov | Difference in strength of regulation of activator and repressor binding to overlapping sites | |
| Avg_corstr_actrep_ov | Average expression correlation of activator and repressor binding to overlapping sites with the target gene | |
| Df_corstr_actrep_ov | Difference in expression correlation of activator and repressor binding to overlapping sites with the target gene | |
| ConsensClustNum | Number of consensus clusters of transcription start sites | Lu and Lin, Genome Res. 2019 |
| ClosestTSS | Closest transcription start site to the coding region | |
| SpreadTSS | Spread of potential transcription start sites | |
| MedPromShapeScore | Median promoter shape score | |
| NumCoopTF_Yang2010 | Number of cooperatively binding regulatory TFs from Yang et al., 2010 data | Yang et al., Cell Research 2010 |
| PercCoopTF_Yang2010 | Percentage of cooperatively binding regulatory TFs from Yang et al., 2010 data | |
| PercOvNocpTF_Yang2010 | Percentage of TFs showing binding site overlaps that are not cooperatively binding TFs as per Yang et al., 2010 data | |
| NumCoopTF_Chen2012 | Number of cooperatively binding regulatory TFs from Chen et al., 2012 data | Chen et al., Bioinformatics 2012 |
| PercCoopTF_Chen2012 | Percentage of cooperatively binding regulatory TFs from Chen et al., 2012 data | |
| PercOvNocpTF_Chen2012 | Percentage of TFs showing binding site overlaps that are not cooperatively binding TFs as per Chen et al., 2010 data | |
| tAI_full | tRNA adaptation index for the full gene | Tuller et al., Cell 2010 |
| tAI_f5 | tRNA adaptation index for the first 5 codons | |
| tAI_f10 | tRNA adaptation index for the first 10 codons | |
| tAI_f15 | tRNA adaptation index for the first 15 codons | |
| tAI_f20 | tRNA adaptation index for the first 20 codons | |
| tAI_f25 | tRNA adaptation index for the first 25 codons | |
| tAI_f30 | tRNA adaptation index for the first 30 codons | |
| tAI_f40 | tRNA adaptation index for the first 40 codons | |
| tAI_f50 | tRNA adaptation index for the first 50 codons | |
| NumGb_H3 | Number of sites in the genebody occupied by H3 | Pokholok et al., Cell 2005 |
| Gb_H3 | Level of H3 in the genebody | |
| NumGb_H4 | Number of sites in the genebody occupied by H4 | |
| Gb_H4 | Level of H4 in the genebody | |
| NumGb_H3K9ac_vsH3 | Number of sites in the genebody showing H3K9ac modification | |
| Gb_H3K9ac_vsH3 | Level of H3K9ac modification in the genebody | |
| NumGb_H3K14ac_vsH3 | Number of sites in the genebody showing H3K14ac modification | |
| Gb_H3K14ac_vsH3 | Level of H3K14ac modification in the genebody | |
| NumGb_H4ac_vsH3 | Number of sites in the genebody showing H4ac modification | |
| Gb_H4ac_vsH3 | Level of H4ac modification in the genebody | |
| NumGb_H3K4me1_vsH3 | Number of sites in the genebody showing H3K4me1 modification | |
| Gb_H3K4me1_vsH3 | Level of H3K4me1 modification in the genebody | |
| NumGb_H3K4me2_vsH3 | Number of sites in the genebody showing H3K4me2 modification | |
| Gb_H3K4me2_vsH3 | Level of H3K4me2 modification in the genebody | |
| NumGb_H3K4me3_vsH3 | Number of sites in the genebody showing H3K4me3 modification | |
| Gb_H3K4me3_vsH3 | Level of H3K4me3 modification in the genebody | |
| NumGb_H3K36me3_vsH3 | Number of sites in the genebody showing H3K36me3 modification | |
| Gb_H3K36me3_vsH3 | Level of H3K36me3 modification in the genebody | |
| NumGb_H3K79me3_vsH3 | Number of sites in the genebody showing H3K79me3 modification | |
| Gb_H3K79me3_vsH3 | Level of H3K79me3 modification in the genebody | |
| NumGb_ESA1 | Number of sites in the genebody showing ESA binding | |

| | | |
|---|---|---|
| Gb_ESA1 | Level of ESA1 binding in the genebody | |
| NumGb_GCN5 | Number of sites in the genebody showing GCN5 binding | |
| Gb_GCN5 | Level of GCN5 binding in the genebody | |
| NumGb_GCN4.AA | Number of sites in the genebody showing GCN4 binding | |
| Gb_GCN4.AA | Level of GCN4 binding in the genebody | |
| NumProm_H3 | Number of sites in the promoter occupied by H3 | |
| Prom_H3 | Level of H3 in the promoter | |
| NumProm_H4 | Number of sites in the promoter occupied by H4 | |
| Prom_H4 | Level of H4 in the promoter | |
| NumProm_H3K9ac_vsH3 | Number of sites in the promoter showing H3K9ac modification | |
| Prom_H3K9ac_vsH3 | Level of H3K9ac modification in the promoter | |
| NumProm_H3K14ac_vsH3 | Number of sites in the promoter showing H3K14ac modification | |
| Prom_H3K14ac_vsH3 | Level of H3K14ac modification in the promoter | |
| NumProm_H4ac_vsH3 | Number of sites in the promoter showing H4ac modification | |
| Prom_H4ac_vsH3 | Level of H4ac modification in the promoter | |
| NumProm_H3K4me1_vsH3 | Number of sites in the promoter showing H3K4me1 modification | |
| Prom_H3K4me1_vsH3 | Level of H3K4me1 modification in the promoter | |
| NumProm_H3K4me2_vsH3 | Number of sites in the promoter showing H3K4me2 modification | |
| Prom_H3K4me2_vsH3 | Level of H3K4me2 modification in the promoter | |
| NumProm_H3K4me3_vsH3 | Number of sites in the promoter showing H3K4me3 modification | |
| Prom_H3K4me3_vsH3 | Level of H3K4me3 modification in the promoter | |
| NumProm_H3K36me3_vsH3 | Number of sites in the promoter showing H3K36me3 modification | |
| Prom_H3K36me3_vsH3 | Level of H3K36me3 modification in the promoter | |
| NumProm_H3K79me3_vsH3 | Number of sites in the promoter showing H3K79me3 modification | |
| Prom_H3K79me3_vsH3 | Level of H3K79me3 modification in the promoter | |
| NumProm_ESA1 | Number of sites in the promoter showing ESA1 binding | |
| Prom_ESA1 | Level of ESA1 binding in the promoter | |
| NumProm_GCN5 | Number of sites in the promoter showing GCN5 binding | |
| Prom_GCN5 | Level of GCN5 binding in the promoter | |
| NumProm_GCN4.AA | Number of sites in the promoter showing GCN4 binding | |
| Prom_GCN4.AA | Level of GCN4 binding in the promoter | |
| Intra_Gb_NumInt | Number of intra-chromosomal interactions in the genebody | Duan et al;, Nature 2010 |
| Intra_Prom_NumInt | Number of intra-chromosomal interactions in the romoter | |
| Inter_Gb_NumInt | Number of inter-chromosomal interactions in the genebody | |
| Inter_Prom_NumInt | Number of inter-chromosomal interactions in the romoter | |
| mRNA_PARS1 | mRNA secondary structure PARS score of the first codon | Kertesz et al., Nature 2010 |
| mRNA_PARS3 | mRNA secondary structure PARS score of the first three codons | |
| mRNA_PARS5 | mRNA secondary structure PARS score of the first five codons | |
| mRNA_PARS10 | mRNA secondary structure PARS score of the first ten codons | |
| mRNA_PARS15 | mRNA secondary structure PARS score of the first fifteen codons | |
| mRNA_PARS20 | mRNA secondary structure PARS score of the first twenty codons | |
| mRNA_PARS25 | mRNA secondary structure PARS score of the first twenty-five codons | |
| mRNA_PARS50 | mRNA secondary structure PARS score of the first fifty codons | |
| mRNA_HL_Mins | mRNA half-life in minutes | Geisberg et al., Cell 2014 |
| protein_HL_Mins | protein half-life in minutes | Belle et al., PNAS 2006 |
| Sun2012_mRNA_Synth_rate | mRNA synthesis rate | Sun et al., Genome Res. 2012 |
| Sun2012_mRNA_Decay_rate | mRNA decay rate | |
| Pugh2004_SAGA_Dominance | Genes showing SAGA dominance in the promoter (Yes/No) | Huisinga and Pugh, Mol. Cell 2004 |
| Pugh2004_TFIID_Dominance | Genes showing TFIID Dominance in the promoter (Yes/No) | |

| | | |
|---|---|---|
| Pugh2004_SAGA_TFIID | Genes activated by both SAGA/TFIID complexes (Yes/No) | |
| Donczew2020_Coactivator_redundant_motif | Number of the coactivator redundant motif present in the promoter | Donczew et al., eLife 2020 |
| Donczew2020_TFIID_dependent_motif | Number of the TFIID redundant motif present in the promoter | |
| TBP.NSMB2009_pol_II | Whether the promoter is a polII transcribed promoter (Yes/No) | van Werven et al., Nat. Struct. Mol. Biol 2009 |
| TBP.NSMB2009_pol_III | Whether the promoter is a polIII transcribed promoter (Yes/No) | |
| TBP.NSMB2009_0 | Ratio of inducible TBP expression level to constitutively expressed TBP level at time t=0 | |
| TBP.NSMB2009_10 | Ratio of inducible TBP expression level to constitutively expressed TBP level at time t=10 mins | |
| TBP.NSMB2009_20 | Ratio of inducible TBP expression level to constitutively expressed TBP level at time t=20 mins | |
| TBP.NSMB2009_25 | Ratio of inducible TBP expression level to constitutively expressed TBP level at time t=25 mins | |
| TBP.NSMB2009_30 | Ratio of inducible TBP expression level to constitutively expressed TBP level at time t=30 mins | |
| TBP.NSMB2009_40 | Ratio of inducible TBP expression level to constitutively expressed TBP level at time t=40 mins | |
| TBP.NSMB2009_60 | Ratio of inducible TBP expression level to constitutively expressed TBP level at time t=60 mins | |
| TBP.NSMB2009_90 | Ratio of inducible TBP expression level to constitutively expressed TBP level at time t=90 mins | |
| TBP.NSMB2009_TBP_occupancy | Overall TBP occupancy | |
| TBP.NSMB2009_TBP_turnover | TBP turnover rate | |
| HolstegeMSB2020_Abf1_NormBindingt0_Gene | Normalized binding levels of Abf1 in the coding region before nuclear depletion of Abf1 | de Jonge et al., Mol. Syst. Biol. 2020 |
| HolstegeMSB2020_Abf1_EstBindingt0_Gene | Estimate for binding levels of Abf1 in the coding region before nuclear depletion of Abf1 | |
| HolstegeMSB2020_Abf1_Offrate_Gene | Abf1 binding offrate in the coding region | |
| HolstegeMSB2020_Abf1_MeanRestimeMins_Gene | Mean residence time of Abf1 in mins in the coding region | |
| HolstegeMSB2020_Abf1_NormBindingt0_Prom | Normalized binding levels of Abf1 in the promoter region before nuclear depletion of Abf1 | |
| HolstegeMSB2020_Abf1_EstBindingt0_Prom | Estimate for binding levels of Abf1 in the promoter region before nuclear depletion of Abf1 | |
| HolstegeMSB2020_Abf1_Offrate_Prom | Abf1 binding offrate in the promoter region | |
| HolstegeMSB2020_Abf1_MeanRestimeMins_Prom | Mean residence time of Abf1 in mins in the promoter region | |
| MolClutch_Nature2012_Rap1Residency_Gene | Residency of Rap1 in the coding region (in mins) | Lickwar et al., Nature 2012 |
| MolClutch_Nature2012_Rap1Occupancy_Gene | Occupancy of Rap1 in the coding region | |
| MolClutch_Nature2012_Rap1Residency_Prom | Residency of Rap1 in the promoter region (in mins) | |
| MolClutch_Nature2012_Rap1Occupancy_Prom | Occupancy of Rap1 in the promoter region | |
| GSE44200_2.5MNase_TBPocc_Gene | TBP occupancy in the coding region (results from 2.5min Mnase treatment) | Zentner and Henikoff, Molecular and Cellular Biology, 2013 |
| GSE44200_2.5MNase_Mot1occ_Gene | Mot1 occupancy in the coding region (results from 2.5min Mnase treatment) | |
| GSE44200_2.5MNase_Mot1TBPrat_Gene | Ratio of Mot1 to TBP occupancy in the coding region (results from 2.5min Mnase treatment) | |

| | | |
|---|---|---|
| GSE44200_2.5MNase_TBPocc_Prom | TBP occupancy in the promoter region (results from 2.5min Mnase treatment) | |
| GSE44200_2.5MNase_Mot1occ_Prom | Mot1 occupancy in the promoter region (results from 2.5min Mnase treatment) | |
| GSE44200_2.5MNase_Mot1TBPrat_Prom | Ratio of Mot1 to TBP occupancy in the promoter region (results from 2.5min Mnase treatment) | |
| GSE44200_10MNase_TBPocc_Gene | TBP occupancy in the coding region (results from 10min Mnase treatment) | |
| GSE44200_10MNase_Mot1occ_Gene | Mot1 occupancy in the coding region (results from 10min Mnase treatment) | |
| GSE44200_10MNase_Mot1TBPrat_Gene | Ratio of Mot1 to TBP occupancy in the coding region (results from 10min Mnase treatment) | |
| GSE44200_10MNase_TBPocc_Prom | TBP occupancy in the promoter region (results from 10min Mnase treatment) | |
| GSE44200_10MNase_Mot1occ_Prom | Mot1 occupancy in the promoter region (results from 10min Mnase treatment) | |
| GSE44200_10MNase_Mot1TBPrat_Prom | Ratio of Mot1 to TBP occupancy in the promoter region (results from 10min Mnase treatment) | |
| GSE59523_NucleosomeAsymmetry_Prom | Nucleosome assymetry (+1 or -1) in the promoter | Ramachandran et al., Genome Res. 2015 |
| YenEtAl_Cell2012_Arp5_Nuc | Whether the gene has Arp5 bound nucelosome (Yes/No) | Yen et al., Cell 2012 |
| YenEtAl_Cell2012_Ino80_Nuc | Whether the gene has Ino80 bound nucelosome (Yes/No) | |
| YenEtAl_Cell2012_Ioc3_Nuc | Whether the gene has Ioc3 bound nucelosome (Yes/No) | |
| YenEtAl_Cell2012_Ioc4_Nuc | Whether the gene has Ioc4 bound nucelosome (Yes/No) | |
| YenEtAl_Cell2012_Isw1_Nuc | Whether the gene has Isw1 bound nucelosome (Yes/No) | |
| YenEtAl_Cell2012_Isw2_Nuc | Whether the gene has Isw2 bound nucelosome (Yes/No) | |
| YenEtAl_Cell2012_Rsc8_Nuc | Whether the gene has Rsc8 bound nucelosome (Yes/No) | |
| YenEtAl_Cell2012_Snf2_Nuc | Whether the gene has Snf2 bound nucelosome (Yes/No) | |
| YenEtAl_Cell2012_Ioc4_terminalNuc | Whether the gene has Ioc4 bound terminal nucelosome (Yes/No) | |
| YenEtAl_Cell2012_Ioc3_terminalNuc | Whether the gene has Ioc3 bound terminal nucelosome (Yes/No) | |
| YenEtAl_Cell2012_Ino80_terminalNuc | Whether the gene has Ino80 bound terminal nucelosome (Yes/No) | |
| YenEtAl_Cell2012_Isw1_terminalNuc | Whether the gene has Isw1 bound terminal nucelosome (Yes/No) | |
| YenEtAl_Cell2012_Isw2_terminalNuc | Whether the gene has Isw2 bound terminal nucelosome (Yes/No) | |
| DionScience2007_Gene_G1Lambda | H3 turnover rate in the coding region of G1 arrested yeast | Dion et al., Science 2007 |
| DionScience2007_Gene_G1Lambda_Zscore | H3 turnover rate in the coding region of G1 arrested yeast (Z-score calculated) | |
| DionScience2007_Prom_G1Lambda | H3 turnover rate in the promoter region of G1 arrested yeast | |
| DionScience2007_Prom_G1Lambda_Zscore | H3 turnover rate in the promoter region of G1 arrested yeast (Z-score calculated) | |
| DionScience2007_Gene_H3Occ | H3 occupancy in the coding region | |
| DionScience2007_Gene_NucOcc | Nucleosome occupancy in the coding region | |
| DionScience2007_Prom_H3Occ | H3 occupancy in the promoter region | |
| DionScience2007_Prom_NucOcc | Nucleosome occupancy in the promoter region | |
| DionScience2007_Gene_PolII_t0 | RNA pol II occupancy in the coding region at t=0 | |
| DionScience2007_Gene_PolII_t60 | RNA pol II occupancy in the coding region at t=60 mins | |
| DionScience2007_Prom_PolIII_t0 | RNA pol II occupancy in the promoter region at t=0 | |

| | | |
|---|---|---|
| DionScience2007_Prom_PolIII_t60 | RNA pol II occupancy in the promoter region at t=60 mins | |
| Phosphorylation | Number of residues in the protein with phosphorylation | Ledesma et al., Database 2018 |
| Methylation | Number of residues in the protein with methylation | |
| Acetylation | Number of residues in the protein with acetylation | |
| Ubiquitination | Number of residues in the protein with ubiquitination | |
| Succinylation | Number of residues in the protein with succinylation | |
| Oxidation | Number of residues in the protein showing oxidation | |
| Nitration | Number of residues in the protein showing nitration | |
| NtAcetylation | Number of residues in the protein with N-terminal acrtylation | |
| Glycosylation | Number of residues in the protein with glycosylation | |
| Ca | Number of calcium binding sites in the protein | |
| Disulfide | Number of residues in the protein showing disulfide bond formation | |
| Lipidation | Number of residues in the protein with lipidation | |
| ActiveSite | Number of residues in the active site of the protein | |
| Sumoylation | Number of residues in the protein with SUMOylation | |
